# Identification of a new inhibitor of Ran GTPase with potential therapeutic value in epithelial ovarian cancer

**DOI:** 10.1101/2025.03.02.641080

**Authors:** Zied Boudhraa, Xiaohong Tian, Ryhem Gam, Sabrina Ritch, Jennifer Kendall-Dupont, Euridice Carmona, Diane Provencher, Jian Hui Wu, Anne-Marie Mes-Masson

## Abstract

Compiled research studies suggest that the small GTPase Ran is critical in cancer initiation and progression. Compounds that block Ran activity would be valuable for cancer treatment. Yet, to date, there is no inhibitor proven to be efficient and specific to Ran. Here, by learning lessons from the discovery of KRAS G12C inhibitors, we generated a structural model of the switch II pocket of GDP-bound Ran to identify potential inhibitors of Ran by virtual screening. After *in vitro* verification, compound M26 was detected. Subsequent hit optimization lead us to identify compound M36 as an inhibitor of Ran with a promising therapeutic value. Binding of M36 to Ran was confirmed by cellular thermal shift assay. The specificity of M36 towards Ran was demonstrated by the evaluation of the active GTP-bound forms of a range of GTPases including Ran, RhoA, Cdc42 and Rac1; and by the expression of a dominant active mutant of Ran. Remarkably, similar to depletion of Ran by siRNA, M36 exhibits a specific toxicity in aneuploid ovarian cancer cells and represses DNA repair systems. In accordance with this, we demonstrated a synergistic relationship between M36 and the FDA approved PARP inhibitor Olaparib. *In vivo*, M36 presents acceptable pharmacokinetic properties and, more importantly, inhibits the tumor growth of an aggressive epithelial ovarian cancer xenograft model. Clinically relevant, M36 was able to induce cell death in *ex vivo* EOC patient derived micro-dissected tumor. Overall, our study is the first to provide a small-molecule compound inhibitor of Ran with a promising therapeutic potential.

## INTRODUCTION

Ran (Ras related nuclear protein), a member of the RAS superfamily of small GTPases, is unique among other GTPases owing to its cellular localization. While other GTPases are found in the cytoplasm and associated to cellular membranes, Ran is found predominately in the nucleus ^1^. Ran has been widely studied for its role in mitosis and nucleo-cytoplasmic shuttling of proteins and RNAs ^2^. In comparison with normal tissues, Ran was found to be overexpressed in several cancers including breast, renal, gastric, colon, pancreatic, ovarian and lung cancers ^3–10^. Furthermore, within the same cancer, the expression of Ran has been found to be positively correlated with patient outcomes in myeloma, lymphoma, neuroblastoma, renal cell, ovarian and breast carcinomas ^4–6,10–14^. It has been reported that Ran is involved in supporting at least three hallmarks of cancer, namely proliferative signaling, resisting cell death and activating pathways that support invasion and metastasis ^7,9,11,15–17^. Interestingly, we and others have been reported that the loss of Ran using specific siRNAs is lethal for cancer cells with no or moderate effect in normal cells ^7,18–21^. This suggests that targeting Ran could achieve a valuable therapeutic index.

Epithelial ovarian cancer (EOC) is the most common form of all ovarian cancers and is the most fatal of all gynecological malignancies ^22,23^. Histologically, EOCs are subdivided into five subtypes where the high-grade serous carcinoma (HGSC) is overrepresented and rates among the most aggressive ^22,24^. EOCs are characterized by extremely high intratumoral heterogeneity that poses specific challenges for therapeutic strategies ^24,25^. However, despite this heterogeneity, aneuploidy is a hallmark of EOC, particularly of HGSC ^26^ where the chromosomal stability control gene, *TP53*, is mutated in >90% of cases ^27,28^. Thus, targeting aneuploidy could potentially treat this cancer.

Recently, we have shown, using EOC models, that the differential sensitivity between normal and cancer cells following the loss of Ran is governed by aneuploidy and genomic instability ^19^. In particular, we have reported that induction of aneuploidy in rare diploid EOC cell lines or normal cells renders them highly dependent on Ran. We also established an inverse correlation between Ran and the tumor suppressor NR1D1 and reveal the critical role of Ran/NR1D1 axis in aneuploidy-associated endogenous DNA damage repair. We showed that Ran, through the maturation of miR4472, destabilizes the mRNA of NR1D1 impacting several DNA repair pathways ^19^. Loss of Ran was then associated with NR1D1 induction, accumulation of DNA damages and lethality of aneuploid EOC cells. Our findings suggest a synthetic lethal strategy targeting aneuploid cells based on their dependency to Ran. We hypothesize that developing small molecules targeting this GTPase is an attractive therapeutic approach for EOC. To date, despite the availability of Ran’s crystal structure ^29–33^, there is no inhibitor proven to be efficient and specific to Ran. The high affinity of Ran to its cognate GTP (pM range) and the high cellular concentration of GTP (millimolar range), makes the identification of an efficient compound with a GTP-competitive mode of action very difficult, if not impossible.

In the present study, inspired by the ground-breaking work of KRAS G12C inhibitors ^34,35^, we aimed to identify allosteric small molecule inhibitors of Ran. The approach chosen was to target the GDP-bound form of Ran at the switch II pocket, with the hypothesis that this would lock the protein in an inactive state, thereby depleting the active Ran population. To this end, a part of the NCI chemical database (90,000 compounds) was screened virtually. After *in vitro* validation and hit optimization, we identified M36 as a promising inhibitor of Ran. Binding of M36 to Ran was confirmed by cellular thermal shift assay. M36 mimics the effect of Ran knock down (KD): specific toxicity in aneuploid cancer cells and significant repression of the DNA damage repair (DDR) process. *In vivo*, M36 is well tolerated by mice at high doses, displays an acceptable pharmacokinetic (PK) pattern and significantly delays tumor growth of a very aggressive EOC xenograft model. At the clinical level, M36 was able to induce cell death in *ex vivo* micro-dissected tumors derived from an EOC patient. Furthermore, owing to its ability to repress DNA repair systems, we established a synergistic relationship between M36 and the FDA approved PARP inhibitor Olaparib. Overall, our findings suggest a new therapeutic strategy for EOC based on the modulation of Ran in aneuploid cells.

## RESULTS

### *In silico* and cell-based screenings identified M26 as hit compound for Ran inhibition

Inspired by the ground-breaking work of KRAS G12C inhibitors, which bind KRAS G12C mutant at a pocket nearby switch II (referred to as switch II pocket) ^34,35^, we aimed to target the GDP-bound form of Ran (RanGDP) at the switch II pocket to ‘arrest’ Ran at its inactivated state (**Figure 1A**). However, one key feature of the switch II pocket of KRAS G12C is that it exists only when it binds with an inhibitor. Our inspection of the crystal structure of RanGDP (pdb: 3ch5) found no pre-formed switch II pocket. We thus decided to construct a structural model of the switch II pocket of RanGDP using the crystal structure of KRAS G12C/Inhibitor complex (pdb: 5v6s) as the template **(Figure 1A)**. Next, a part of the NCI chemical database (90,000 compounds) was screened virtually against the RanGDP switch II pocket. Top-ranking compounds identified in this *in silico* screen went through a more in-depth visual inspection for binding modes. A total of 28 compounds were selected for further study as potential Ran inhibitors (**Figures 1B and S1A**). Having previously established that aneuploid EOC cells are more sensitive to the loss of Ran than normal or tumoral diploid cells ^19^, biological activity was assessed by clonogenic assays (at a single dose of 10 µM) using one aneuploid EOC cell line (TOV112D) and the diploid normal ARPE (ARPE-19) cells **(Figures 1C and S1B)**. The criterion for a positive hit was that the compound did not inhibit the colony formation of the ARPE cells but significantly inhibited the number of colonies formed by the TOV112D cells. Our results show that one compound from the *in silico* screening, M26 **(Figure 1B)**, inhibited colony formation of EOC but not the normal cells **(Figure 1C)**. Furthermore, by analyzing PARP cleavage by Western blot, we found that M26 treatment induced apoptosis in TOV112D cells with no notable effect in ARPE cells **(Figure 1D)**, indicating a differential sensitivity between the diploid and aneuploid cells following M26 treatment. Since our screening was virtual using Ran crystal structure, it was therefore important to determine whether our hit compound was able to inhibit Ran in its biological form. For that, the level of active GTP-bound Ran was evaluated in lysates of EOC cells treated or not with M26 using Ran pull down assays. Because of the fast turnover between Ran-GTP and Ran-GDP, a fast incubation time (1 h) and a higher compound concentration is recommended for this pull down assay, as described in previous publications regarding other GTPases, such as RalA ^36^. Our results showed a marked decrease in Ran-GTP level in TOV112D cells incubated with M26 compared to DMSO-treated control cells **(Figure 1E)**. Collectively, our data identified M26 as a potential inhibitor of Ran with an interesting therapeutic value.

**Figure 1:**
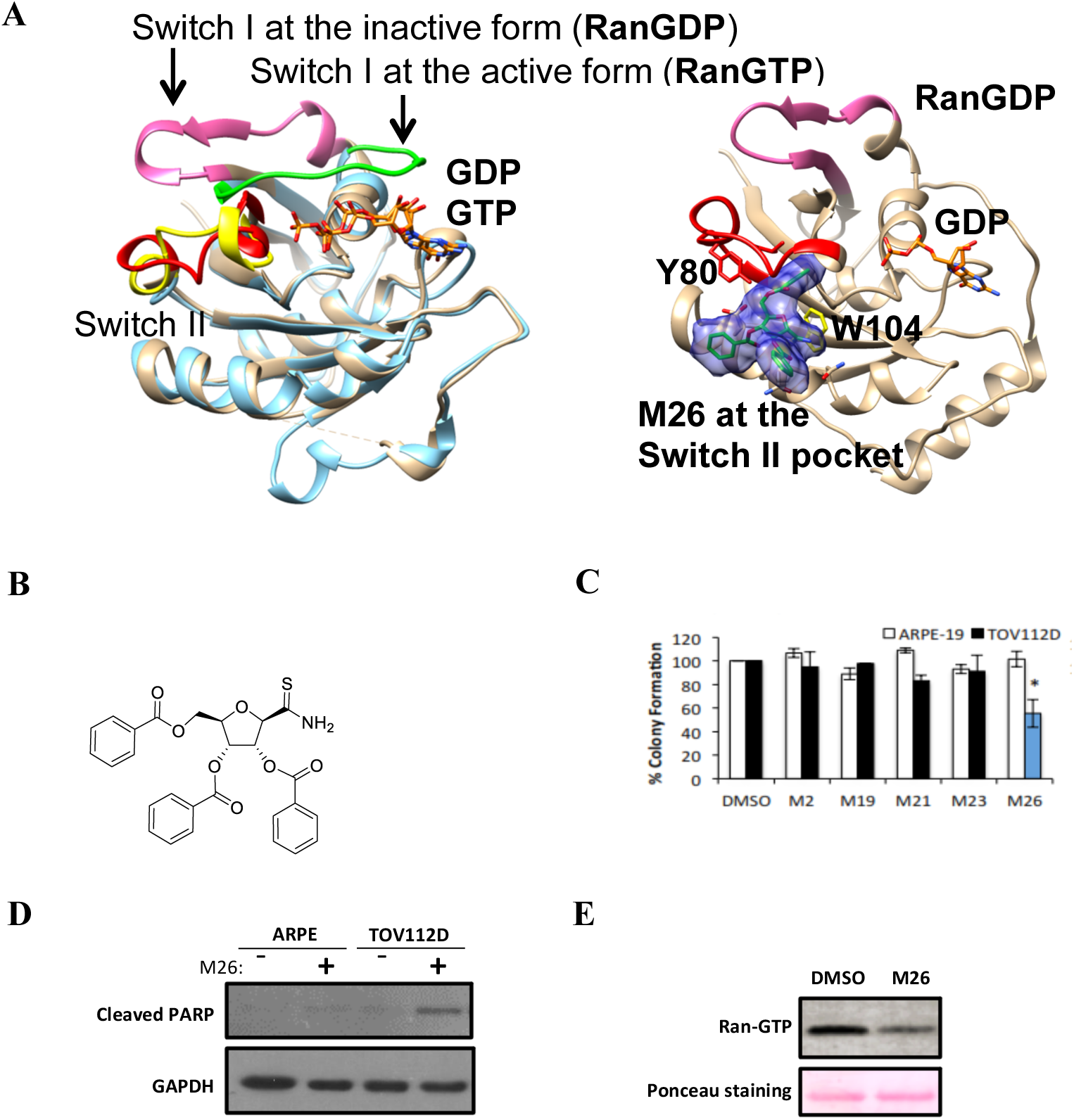
*In silico* and cell-based screenings identified M26 as a hit compound for Ran inhibition. **(A)** Left side: Crystal structure of RanGDP (pdb: 3ch5) was aligned to RanGTP (pdb: 3nc1), showing different conformations of Switch I at the activated and inactivated states; Right side: Structural model of the switch II pocket of RanGDP was built by homology modeling and the predicted binding mode of M26 (green sticks with blue surface) at the switch II pocket of RanGDP was showed. **(B)** Chemical structure of M26. **(C)** Colony formation inhibition by selected M compounds at 10 µM in normal ARPE and EOC TOV112D cells. Only the compound M26 inhibited TOV112D without affecting ARPE (blue bar). Bars represent percentage of colonies formed compared to DMSO-treated controls. **(D)** ARPE and TOV112D cells were incubated with M26 to evaluate apoptosis by western blot using the antibody anti-cleaved PARP. **(E)** TOV112D cells were incubated for one hour with M26 (50 µM) before performing RanGTP pull-down assay on the resultant cell extract.

### Chemical optimization and cell-based screening identified M36 as a more promising inhibitor of Ran

Plasma esterases are the major drug-metabolizing enzymes in plasma significantly affecting drug stability *in vivo* ^37^. As shown in **Figure 1B**, M26 has three ester groups, which would suggest poor bioavailability of the drug when injected *in vivo*. To overcome this issue and explore preliminary structure-activity relationship, a series of M26-derived compounds, in which the ester groups were replaced by ether groups, were synthetized **(Figures 2A and S2A)**. Our results showed that only compound M36 and its derivative M98, inhibited colony formation of TOV112D cells without affecting the ARPE cells **(Figure S2B)**. Interestingly, M36 but not M33 inhibited the activation of Ran comparably to the parent compound, M26 **(Figure S2C)**. Therefore, for the rest of the study we selected M36 **(Figure 2A)** for further characterization. Using Ran pull down assays, the effect of M36 on Ran activation was shown to be dose dependent **(Figure 2B)**. In addition, immunofluorescence studies using an antibody specific to active Ran-GTP showed no detectable levels of Ran-GTP in TOV112D cells treated with M36 as compared to the intense staining of the DMSO-treated cells **(Figure 2C)**. More importantly, the direct binding of M36 to RanGTPase was assessed by cellular thermal shift assay (CETSA), a widely used technique to verify the interaction of a compound with its target protein ^38^. In this assay, the thermal stability of Ran was investigated at different temperatures in OV1946 cells treated with different concentrations of M36 or with DMSO **(Figures S3A-B)**. Like the pull down assay, shorter incubation time (1 h) and higher compound concentrations are needed for CETSA in order to detect binding of no-covalent interactions. As expected, Ran levels detected by Western-blot decreased with increasing temperatures in controls cells treated with vehicle DMSO. However, a shift in this thermal instability was observed when cells were treated with M36, specially at the concentrations of 100 and 200 µM. Significantly higher levels of Ran were observed at all M36 concentrations for temperatures of 58°C, 59°C and 60°C **(Figure S3A)**, indicating M36 engages with Ran and increases its thermal stability. As negative control, we show that this protection was not observed when OV1946 cells were treated with the same increasing concentrations (50-200 µM) of the PARP inhibitor Olaparib (**Figure S3C**), a drug known to impact DNA repair ^39^. Since this CETSA relies on densitometry analyses of Western blot bands, we confirmed these results with the more quantitative HiBiT CETSA ^40^. This method uses a commercially available luminescent tag (HiBiT from Promega) for protein thermal stability assessment. Here, OV1946 cells were transiently transfected for 24 h with Ran cDNA having N-terminal HiBiT tag (see Methods for details), treated with 100 µM M36 or DMSO for 1 h, and heated for 3 min at different temperatures. Luminescence from the HiBiT tag was then measured in cell lysates. As expected luminescence signal decreased with increasing temperatures, and a shift in this thermal stability was observed when cells were treated with M36 when compared to DMSO treated cells (**Figure 2D**). Significant differences were observed at temperatures from 57°C to 62°C, corroborating the CETSA results using Western blot. Overall, these results indicate that M36 targets Ran and inhibits Ran-GTP activation.

**Figure 2:**
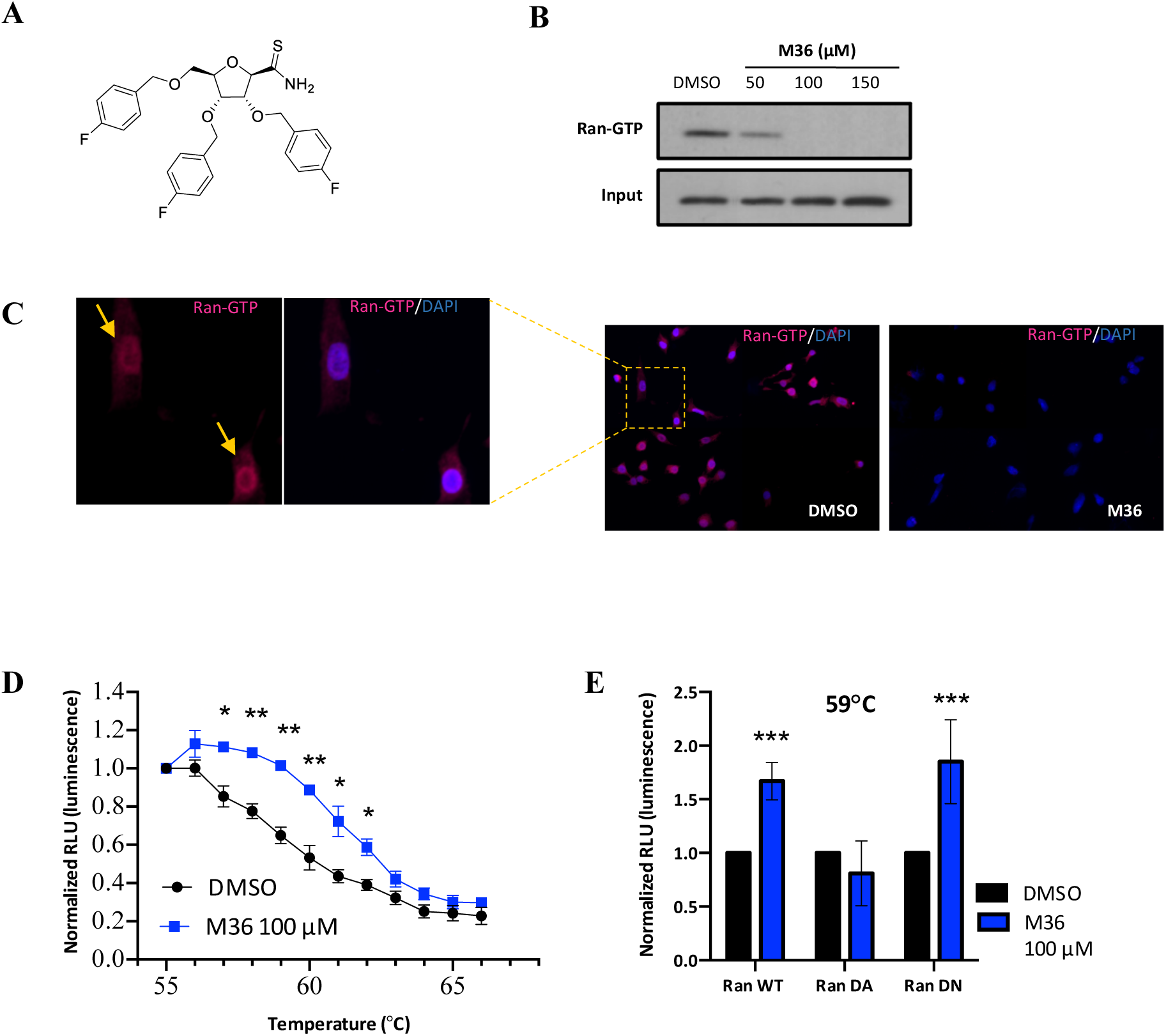
M36, a derivative of M26, binds and impacts Ran activation in EOC cells. **(A)** Chemical structure of M36. **(B)** TOV112D cells were treated for one hour with increasing concentrations of M36. Subsequently, RanGTP pull-down assay was performed to assess Ran activation. **(C)** TOV112D cells were treated with M36 (100 µM) for one hour. After cell fixation and permeabilization, RanGTP was revealed by immunofluorescence using a specific antibody (pink). Nuclei are shown by Dapi staining (blue). **(D)** OV1946 cells were transiently transfected with Ran cDNA containing N-Terminal HiBiT or with empty plasmid, and then treated with 100 µM M36 (blue square symbols) or DMSO (black round symbols) for one hour (see Methods for detail). Luminescence of the HiBiT tag was evaluated in cell extracts after heating the cells at different temperatures for three minutes. Values represent luminescence readings normalized to the background value of the empty plasmid and to the first temperature used (55°C). **(E)** OV1946 cells were transiently transfected with HiBiT Ran WT or mutants dominant active Ran^Q69L^ or dominant negative Ran^T24N^ and then treated with 100 µM M36 (blue bars) or DMSO (black bars) for one hour. Luminescence of the HiBiT tag was evaluated in cell extracts after heating the cells at 59°C for three minutes. Values represent luminescence readings normalized to the DMSO treated cells. *P < 0.05 **P < 0.01 *** P < 0.001 (n = 3, Student’s t-test).

Regarding its effect on cancer cells, the IC_50_ value of M36 for the TOV112D cell line as determined by clonogenic assay was found to be 10.02±0.36 µM. Only a minimal effect was observed for the normal ARPE cell line at high concentrations **(Figure 3A)**. To support our observation regarding the differential sensitivity between diploid and aneuploid cells with M36 treatment, we expanded our analysis by using, in addition to ARPE and TOV112D cell lines, five aneuploid HGSC cell lines [TOV1946, TOV2295(R), OV1946, OV866(2) and OV4485] and one diploid low-grade serous EOC TOV81D cell line. Proliferation assays were performed on these cell lines after treatment with M36 and a concentration higher than the observed IC_50_ value was chosen (40 µM) to better evaluate differential effects. As a positive control, the same cell lines were transfected with siRan that we have previously characterized for its specificity ^19,41^. Remarkably, like siRan, M36 demonstrated a specific toxicity in aneuploid cells with only a moderate effect in normal and EOC diploid cells **(Figure 3B)**. Similarly, this toxicity was accompanied by the induction of apoptosis specifically in aneuploid cells **(Figure 3C)**, highlighting the therapeutic potential of M36. Since aneuploidy and genomic instability are hallmarks of the majority of cancers ^42,43^, we assessed the toxicity of M36 in other cancer cell lines derived from prostate, gastrointestinal, breast and skin cancers. Strikingly, M36 showed a significant toxicity in all the cell lines tested **(Figure S4A)**. These results suggest that the use of M36 would not be limited to ovarian cancer but could be extended to other tumors. To investigate the specificity of M36 towards Ran, the effect of M36 on the activation of other GTPases (RhoA, Cdc42 and Rac1) was assessed. As shown in **Figure 3D**, no decrease in the activation of these GTPases was observed when incubated with M36. In corroboration, no thermal stability of Cdc42 was observed on CETSA after M36 treatment **(Figure S3D)**. Furthermore, overexpression of constitutively active (GTP-bound form) Ran^Q69L^ mutant protein ^44^ mitigated the inhibition of TOV112D cell growth by M36, whereas overexpression of dominant negative Ran^T24A^ mutant ^44^ had similar effect as the wild-type (WT) protein **(Figure 3E)**. Furthermore, HiBiT CETSA experiments performed at 59°C in OV1946 cells show that M36 was able to protect Ran WT and Ran^T24A^, but not Ran^Q69L^, from thermal instability **(Figure 2E)**, suggesting that M36 does not bind to the GTP-bound form of Ran. Together, these data provide evidence of the specificity of M36 towards Ran and indicate that this compound acts specifically on the GDP-bound form of Ran.

**Figure 3:**
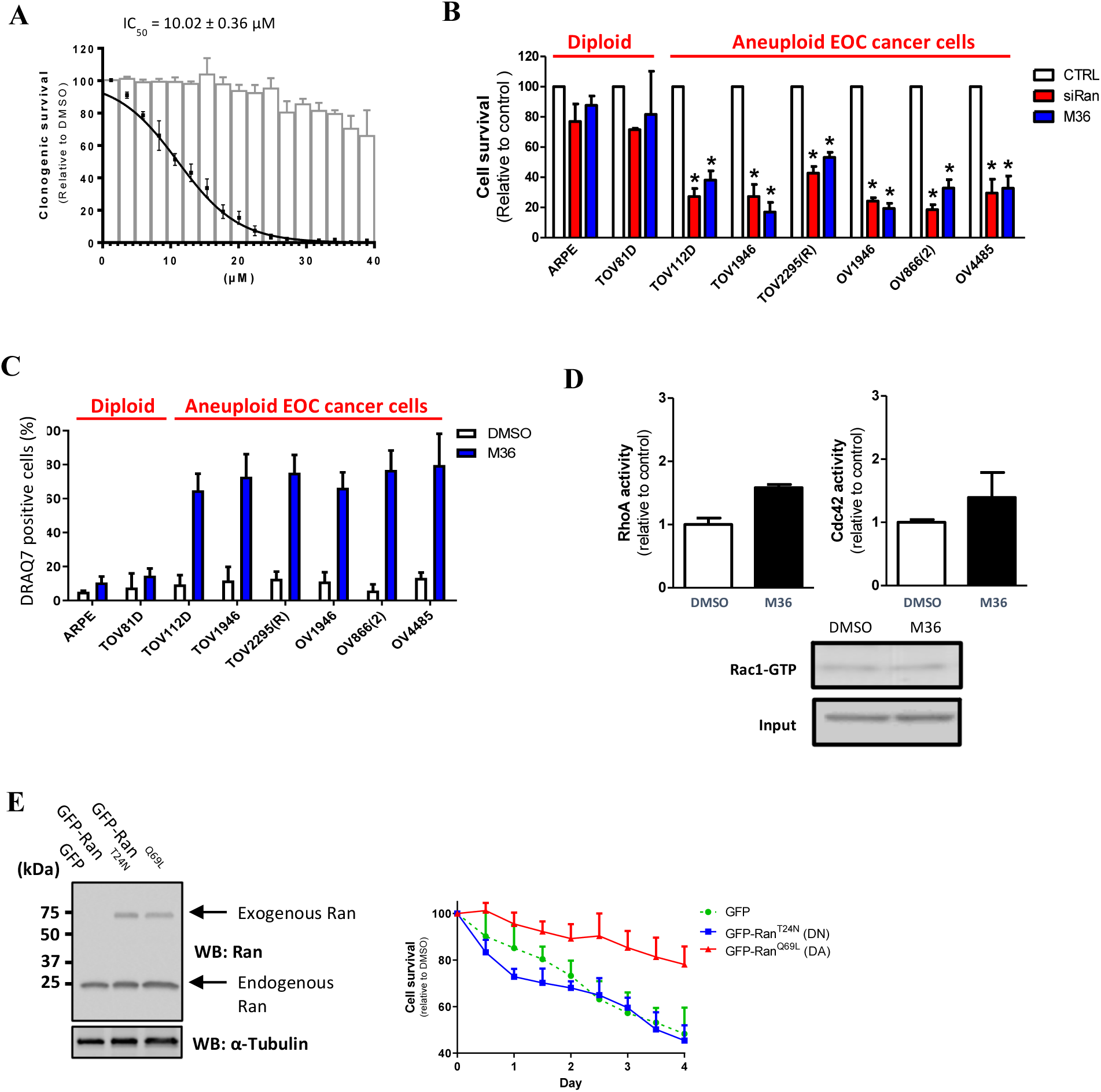
M36 shows selectivity for aneuploid cells and does not interferes with the activation of other GTPases. **(A)** Clonogenic assay using different concentrations of M36 in ARPE and TOV112D cells. IC_50_ value was calculated for the TOV112D cells. Bars and curve points represent percentage of colonies formed compared to DMSO-treated controls in ARPE and TOV112D cells, respectively. **(B)** different EOC cell lines and the normal ARPE cells were transfected with a siRNA targeting Ran or with M36 (40 µM) before assessing cell proliferation using the live cell imaging IncuCyte system. Data are expressed as the percentage of control cells at the end of experiment (96 h) and are representative of at least three independent experiments. **(C)** ARPE and EOC cells were treated with M36 (40 µM) for 6 days and then cell death was assessed by FACS using DRAQ7 staining. **(D)** TOV112D cells were treated for one hour with M36 (50 µM) in order to evaluate RhoA (left), Cdc42 (right) and Rac1 (bottom) activities. **(E)** Immunoblots showing successful overexpression in TOV112D cells of dominant active Ran^Q69L^ and dominant negative Ran^T24N^ (left panel). These cells were treated with M36 and cell proliferation was assessed using the live cell imaging IncuCyte system (right panel). Data are expressed as the percentage of DMSO treated cells. *P < 0.05 (n ≥ 3, Student’s t-test).

### M36 mimics the effect of Ran knock down (KD) regarding DDR

We have shown that the differential sensitivity between diploid and aneuploid cells following the loss of Ran comes from the involvement of the Ran/NR1D1 axis in modulating DNA repair ^19^. In fact, following the loss of Ran, NR1D1 was induced and in turn repressed several DNA repair pathways including the homologous recombination (HR) and the non-homologous end joining (NHEJ) pathways, leading to the accumulation of DNA damages. We first assessed the effect of M36 on NR1D1 expression. As observed following Ran KD, M36 significantly induced the expression of NR1D1 in two aneuploid EOC cell lines (TOV112D and OV1946) **(Figure 4A-B)**. As negative control, we have shown that NR1D1 expression was not affected when cells were treated with the PARP inhibitor Olaparib **(Figure S4B)**, suggesting that M36 effect on NR1D1 expression is specific. Next, to investigate whether M36 was able to repress the DDR process, TOV112D cells were either transfected with siRan or treated with M36 and the DDR components were analyzed by immunofluorescence. Remarkably, in accordance with Ran KD, M36 treatment induced DNA damage accumulation as shown by an increase of p-ɣH2AX foci number **(Figure 4C)** and an inhibition of the HR and the NHEJ pathways, which were assessed respectively by the quantification of Rad51 **(Figure 4D)** and 53BP1 **(Figure 4E)** foci formation. Interestingly, since the cytoplasmic staining of 53BP1 following M36 treatment was increased **(Figure 4F)**, the inhibition of 53BP1 foci formation by M36 can probably be attributed to the inhibition of the Ran-dependent nuclear import of this protein that has been previously reported ^15^ and is in accordance with the Ran KD condition **(Figure 4F)**. Collectively, our data demonstrate that M36 mimics the effect of the loss of Ran regarding DDR.

**Figure 4:**
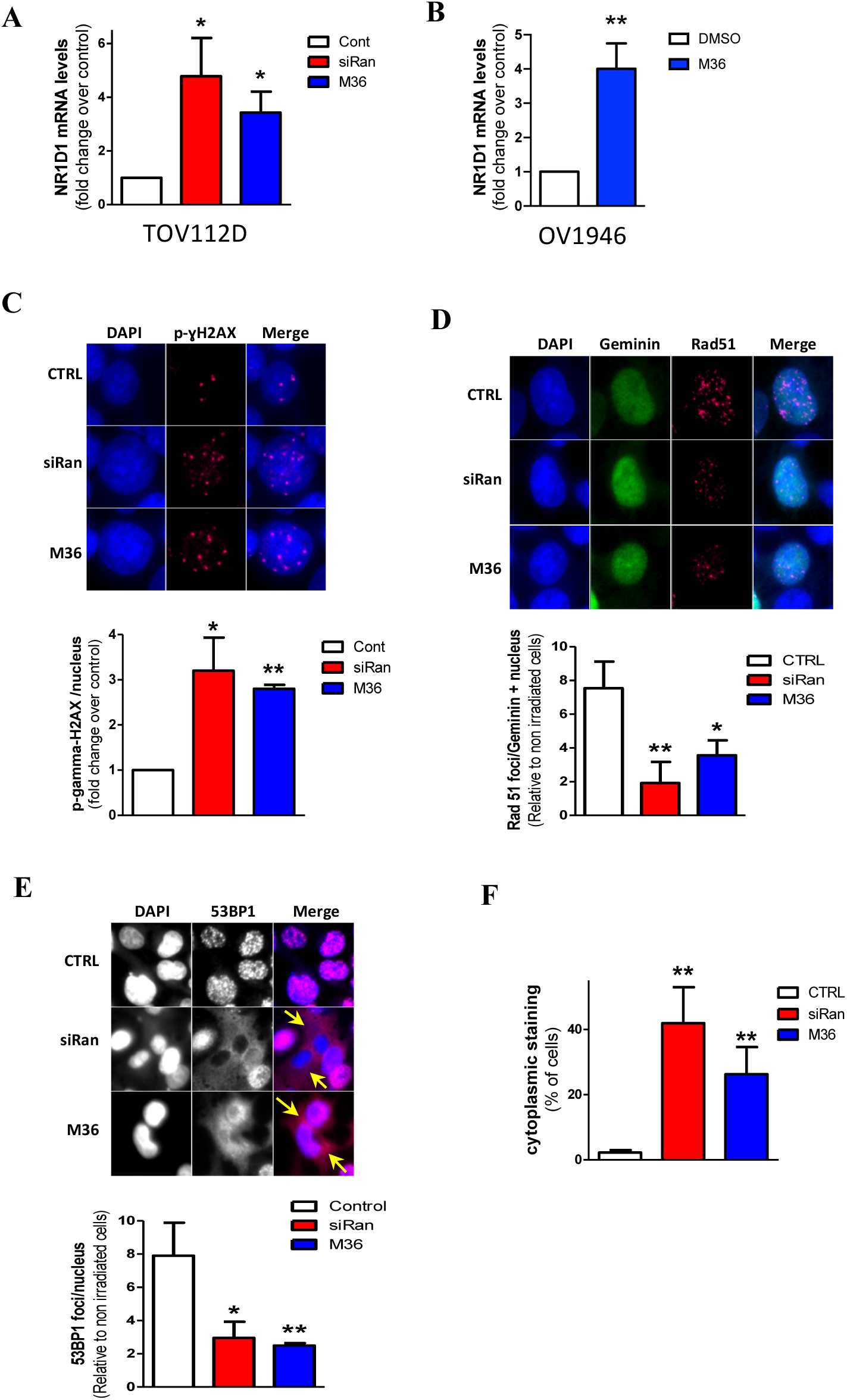
M36 mimics the effect of Ran KD regarding DDR. **(A)** The expression of NR1D1 was evaluated by qRT-PCR in TOV112D cells transfected with siRan or treated with M36 (40 µM). **(B)** OV1946 cells were treated with M36 (50 µM) and expression of NR1D1 was evaluated by qRT-PCR. **(C-E)** TOV112D cells were treated with siRan or M36 (20 µM) and allowed to grow for 72 hours. For p-ɣH2AX staining **(C)**, cells were fixed and stained. For Rad51 **(D)** and 53BP1 **(E)** staining, cells were gamma-irradiated at 10 Gy and fixed 1 hour after irradiation. Representative images after irradiation **(upper panel)** and quantitative analyses of foci formation **(lower panel)** are presented. For Rad51 and 53BP1 foci, data are expressed as the ratio between foci number after and before irradiation. **(F)** Images obtained from **(E)** were used to quantify cells presenting cytoplasmic staining of 53BP1. *P < 0.05 **P < 0.01 (n ≥ 3, Student’s t-test).

### Evaluation of the therapeutic potential of M36 *in vivo*

The promising *in vitro* results of M36 led us to initiate *in vivo* analysis of this small molecule inhibitor of Ran. PK results showed that after a single intraperitoneal (IP) injection (200 mg/kg), plasma levels of M36 was detected at our first time point of 5 minutes, reached a peak at 30 minutes (100 μM), remained at around 10 μM after 6 h and is still detectable at 24 h **(Figure 5A)**. Therefore, M36 displayed an acceptable PK pattern. For the tolerance study, M36 was injected IP daily for a period of three weeks. Our results showed that treated mice continued to gain weight **(Figure 5B)**, suggesting that, even at higher doses (200 mg/kg), M36 was well tolerated. Inhibition of tumor growth was evaluated in the human chemoresistant EOC xenograft model, TOV112D. As we’ve previously shown, this xenograft model showed resistance to carboplatin treatment ^45^. Furthermore, when mice were treated with the maximum tolerated dose of the PARP inhibitor Olaparib **(Figure 5C)**, no significant effect was observed regarding tumor growth. However, M36 treatment significantly delayed tumor growth of this aggressive model **(Figure 5D-E)** and prolonged mouse survival **(Figure 5F)**, highlighting the promising potential of this compound for the treatment of EOCs, particularly for patients with poor prognosis.

**Figure 5:**
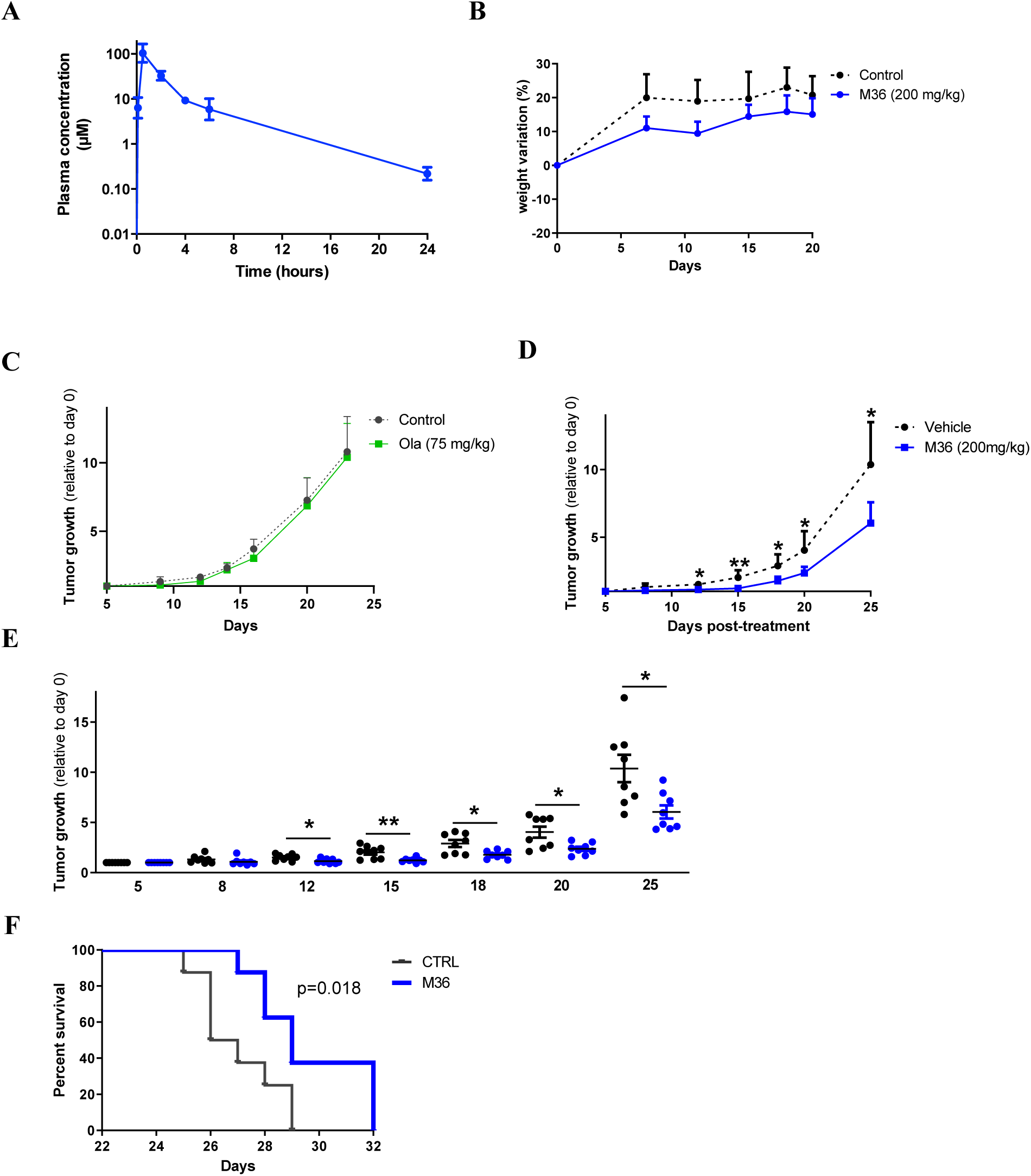
Evaluation of the therapeutic potential of M36 *in vivo*. **(A)** Measurements of plasma levels of M36 compound after a single intraperitoneal injection (200 mg/kg) in CD1 mice. **(B)** M36 (200 mg/kg) was injected IP daily in NRG mice for 3 weeks. Mouse weight was measured at the indicated time points and expressed as the percentage of their weight on the day of the first injection. **(C-D)** TOV112D cells were subcutaneously injected in NRG mice and treatments with Olaparib (75 mg/kg, daily) **(C)**, and M36 (200 mg/kg, daily) **(D)** were performed. Data are normalized with the mean of tumor size at the first day of injection. **(E)** Tumor growth in individual mice treated with M36 is shown. **(F)** Kaplan-Meier survival curve of mice from **(D-E)**. In **(D)** and **(E)** data were analyzed using the two-tail Student’s t test. In **(F)**, data were analyzed using the Log rank test. *P < 0.05, **P < 0.01.

### M36 induces cell death in *ex vivo* ovarian cancer models

To evaluate the clinical relevance of M36 treatment, the viability of microdissected tumor samples derived from EOC xenografts or from a patient tumor were evaluated *ex vivo* using our microfluidic methodology ^46^. Our results show **(Figure 6)** a significant increase in the intensity of cleaved caspase 3 (CC3) staining 72 h after M36 treatment in both the OV1946 xenograft and the TOV12018G patient tumor models. For the TOV112D xenograft model, this effect was also observed, but no significance was achieved **(Figure S4C)**, and a 50% decrease in the nuclear Ki-67 staining was observed in this model 24 h post-treatment. These results suggest that M36 induces cell death by apoptosis and could decrease cell proliferation in *ex vivo* EOC models, including patient tumor, further corroborating its potential therapeutic value.

**Figure 6:**
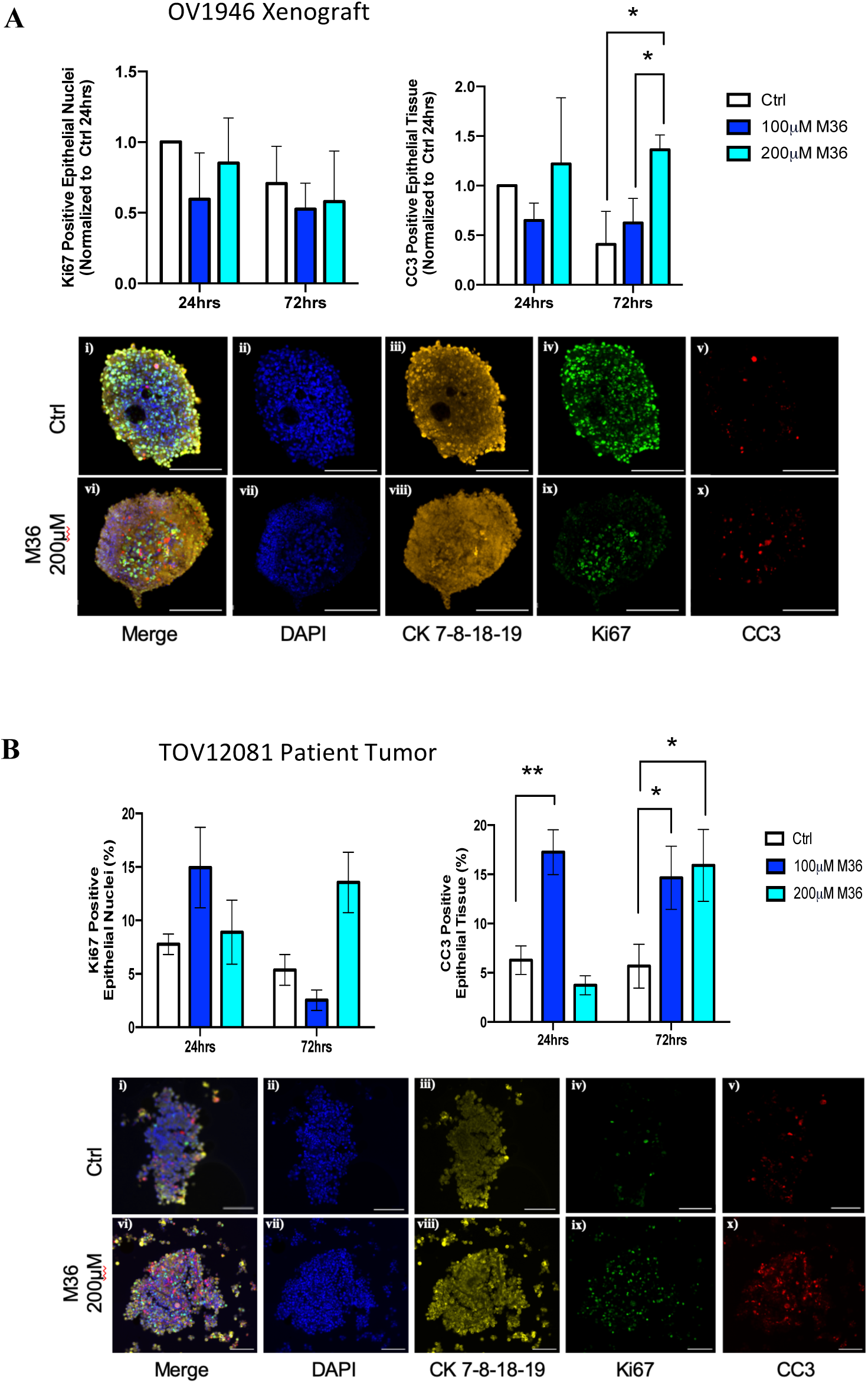
M36 induces cell death of *ex vivo* micro-dissected tumors from xenograft and patient tissue. Samples from OV1946 xenografts **(A)** or TOV12018G patient tumor **(B)** were processed as micro-dissected tissues (MDTs) and cultured in microfluidic devices as described in Methods. MDTs were treated with M36 at different concentrations for 24 or 72 h and then formalin-fixed, paraffin embedded. Sections (6µ) were stained for KI67 (green), cleaved-caspase 3 (red), a cytokeratin cocktail (yellow) and DAPI (blue). Bar graphs represent the expression of nuclear KI67 (left panels) or CC3 (right panels) within the tumor epithelium (CK positive cells) of each MDT. Data shown represents mean ± SEM of three separate experiments, normalized to Ctrl 24hrs. Statistical analysis was done using Two-way ANOVA followed by Tukey’s test. *P < 0.05. Scale bars of representative images are 200 µm.

### Combination of M36 and Olaparib triggers synthetic lethality in EOC cells

It is presumed that resistance to PARP inhibitor therapy is closely related to the activation of the HR pathway of cancer cells ^47^. Therefore, combination treatments of PARP inhibitor with drugs that inhibit the HR pathway would be promising to circumvent this resistance. Based on our finding that M36 was able to repress the HR pathway, we sought to determine whether M36 was able to sensitize EOC cells to Olaparib. First, we investigated the effect of M36 on the activation of the HR pathway by Olaparib. Our data in two HR proficient BRCA1/2 WT EOC cell lines [TOV112D, OV866(2)] showed that M36 pre-treatment efficiently inhibited the induction of the HR pathway (assessed by Rad51 foci quantification) by Olaparib **(Figure 7A)**, and this repression was comparable when irradiation was used as an inducer of this DNA repair pathway **(Figure 4D)**. Next, combination treatments of M36 with Olaparib were undertaken in three BRCA1/2 WT and Olaparib resistant-cell lines, namely TOV112D, OV866(2) and OV90 cell lines, together with the normal diploid ARPE cell line. These combinations were performed using increasing concentrations of both drugs (all concentrations used are below the IC_50_) and evaluated using the Bliss independent model ^48^, in which values below −1 indicate antagonism, values between −1 and 1 indicate additive effects and values above 1 indicate synergy. Remarkably, in contrast to ARPE cells that did not respond to any combination condition, M36 displayed a potent synergistic interaction with Olaparib in all 3 EOC cell lines (Bliss scores ranging from 41.02 to 78.6) **(Figure 7B-D)**. Similarly, while no effect was observed in ARPE cells, M36/Olaparib combination induced a significant increase of cell death in the EOC cell lines in comparison with single treatments, as assessed by FACS using DRAQ7 staining **(Figure 7E)** and by Western blot using cleaved caspase-3 antibody **(Figure 7F)**. Overall, our data indicate that combining M36 with Olaparib would be an effective combination therapy to counter PARP inhibitor resistance in ovarian cancer.

**Figure 7:**
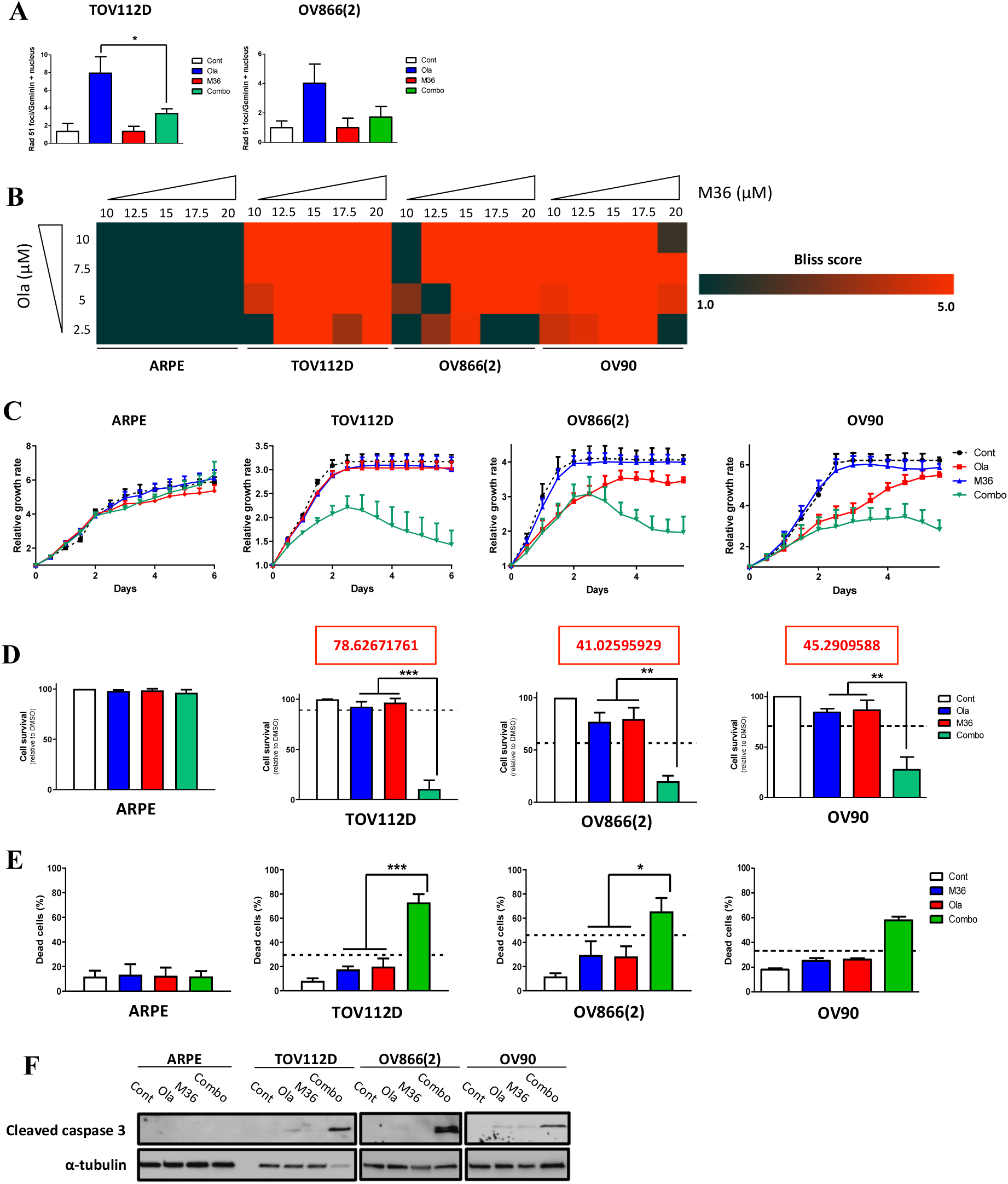
Combination of M36 and Olaparib triggers synthetic lethality in EOC cells. **(A)** TOV112D and OV866(2) cells were treated with M36 (20 µM) for 48 hours then with Olaparib (10 µM) for 24 hours. After fixation, cells were immunostained with Rad51 antibody. Quantitative analyses of foci formation are presented. **(B)** Cells were treated with increasing concentrations of M36 and/or Olaparib. The heat map shows Bliss scores calculated for each combination at day 6. **(C)** For each cell line, an example of proliferation profile following Olaparib and M36 combination treatment is presented. **(D)** At the end of the experiment presented in **(C)**, dead cells were removed by PBS washes and cell confluence was measured using the live cell imaging IncuCyte system. Data are expressed as the percentage of control cells. **(E, F)** ARPE and EOC cells were treated as in **(C)** and apoptosis was assessed by flow cytometry **(E)** and western blot **(F)** using DRAQ7 dye and cleaved caspase 3 antibody, respectively. *P < 0.05 **P < 0.01 *** P < 0.001 (n = 3. Student’s t-test).

In all, we describe here a first-in-class small molecule inhibitor of the Ran GTPase targeting aneuploid EOC. However, further medicinal chemistry is needed to improve potency and compound solubility, and we are aware that M36 is not ultimate inhibitor to go forward to a clinical setting. Using the computational model proposed and the M36 scaffold, we are confident that better compounds targeting the Ran GTPase could be unraveled in a near future.

## DISCUSSION

The involvement of Ran GTPase in cancer initiation and progression has been discussed for more than a decade ^7,8,15,17,21,41,49–51^. Furthermore, using several *in vitro* and *in vivo* experimental models, Ran has been proposed as a promising therapeutic target of cancer ^8,9,14,17–19,52^. Recent reports have proposed new strategies to target Ran in the context of cancer therapy. One of them relies on repurposing Pimozide in Ran Targeting ^53^. Pimozide was initially identified as a potent antagonist of dopamine receptor D2 (D2R) and used as an antipsychotic drug for the management of schizophrenia and other psychotic disorders. It has been shown that Pimozide inhibits the expression of Ran at the transcriptional level and inhibits breast cancer cell growth *in vitro* and *in vivo* ^53^. Although this molecule displays an interesting cancer therapeutic value, it is likely that this drug will display severe side effects at the concentrations needed. In fact, it has been shown that mice lacking D2R had impaired motor activity and movement coordination as well as hyperactivity ^54^. More importantly, the dosage of Pimozide for patients with psychotic disorders is about 0.2 mg/Kg/day, however, in the *in vivo* study proposed for breast cancer therapy, mice were treated daily with Pimozide at 20 mg/Kg/day in order to have a significant effect on tumor growth ^53^. These very high doses of Pimozide will inevitably be an issue in the exploration of this drug in the context of cancer. Another strategy relies on Nano-encapsulation technology (using PLGA-based nanoparticles) of a shRNA targeting Ran ^55^ or a peptide inhibiting the activation of this GTPase by disrupting the interaction between Ran and RCC1 ^56,57^. Such an approach presents many obstacles making its use in clinic very difficult. First, upon entering the circulation, nanoparticles (NPs) are often coated with opsonin proteins leading to their clearance by macrophages and affecting their pharmacokinetic properties ^58^. Second, blood vessels are leaky with fenestrations ranging between 0.2 and 1.2 µm. Since NPs exist in a range of sizes, it is thought that the size of therapeutic NPs should be well below 200 nm ^58,59^. However, the NPs used for Ran targeting have a mean diameter of 182–277 nm ^56^.

Here, we aimed to target Ran directly with a new small molecule inhibitor. To date, there is no available drug specific to Ran. One reason for this is the high intracellular concentration of GTP as well as the high affinity of GTP to its cognate GTPase, making the development of an efficient competitor of GTP difficult. Alternatively, it is thought that developing allosteric compounds able to interact with a given GTPase in its GDP-bound form and to lock it on its inactive state is more relevant to attenuate GTPase activity. Inspired by the work of KRAS G12C inhibitors, which bind KRAS G12C mutant at the switch II pocket ^34,35^, we aimed to target the RanGDP at the switch II pocket to ‘arrest’ Ran at its inactivated state. However, our inspection of the crystal structure of RanGDP (pdb: 3ch5) found no pre-formed switch II pocket. We thus constructed a structural model of RanGDP switch II pocket by homology modeling and performed in silico screening against the switch II pocket. Chemical optimization of our initial hit M26 lead to compound M36 as a novel Ran inhibitor. Importantly, M36 displays, *in vitro,* a very interesting therapeutic index between normal/diploid and cancer/aneuploid cells that is in alignment with our *in vivo* data. In fact, at higher doses, M36 is well tolerated by mice and exhibits a significant toxicity for one of the most aggressive xenograft models, TOV112D, that do not respond to conventional treatments such as Carboplatin or Olaparib therapies. Furthermore, at the clinical level, M36 was able to induce cell death by apoptosis in *ex vivo* micro-dissected tumors derived from an EOC patient. Another advantage of allosteric inhibitors is that they bind to less conserved sites across GTPases and kinases ensuring specificity and avoiding potential severe side effects ^60^. In this regard, we have shown that M36 does not influence the activation of other GTPases such as RhoA, Cdc42 and Rac1. More importantly, overexpression of a dominant active of Ran rescued the toxicity of M36 highlighting the specificity of our Ran inhibitor.

Olaparib has changed the paradigm for HGSC treatment and has been approved by the FDA as maintenance therapy for patients harboring BRCA1/2 mutations (HR deficient) ^61^. This stratification comes from the observation that HR activation is closely linked to non-responsiveness and resistance to PARP inhibitor therapy ^47^. Therefore, combination treatment of a PARP inhibitor with drugs that inhibit the HR pathway is actively being exploited in the clinical setting ^62,63^, and might be effective to induce lethality in HGSC cells. Here, we have shown that M36 induced HR deficiency and dramatically sensitized three BRCA1/2 WT and Olaparib resistant cell lines (TOV112D, OV866(2) and OV90 ^64^) to PARP inhibitor therapy. This result is of great interest since it could open new perspectives to circumvent resistance, and possibly to expand PARP inhibitor therapy to patients that are currently excluded. Many cancers are characterized by widespread aneuploidy and genomic instability ^42,43^. Hence, targeting these hallmarks would be effective for the development of a new targeted therapy, not only for EOC, but also for a variety of cancers. In our previous work, we have shown that targeting Ran is an effective way to eliminate cancer cells with chromosomal and genomic instability ^19^. This prompted us to investigate the therapeutic value of M36 in other cancers. Interestingly, we found that M36 is also toxic in a range of cancer models including prostate, gastrointestinal, breast and skin cancers. These results are encouraging and in future studies we plan to investigate the *in vivo* efficacy of M36 in these models.

In conclusion, though further medicinal chemistry optimization is required, our study is the first to provide the generation of a first-in-class compound inhibitor of Ran with a therapeutic potential for cancer in general and EOC in particular.

## METHODS

### Computational modeling and in silico screening

The structural model of RanGDP with switch II pocket was built using crystal structure of RanGDP (pdb: 3ch5) and crystal structure of KRAS G12C in complex with an inhibitor at switch II pocket (pdb: 5v6s) as the templates, by software Modeller (UCSF, CA, USA). Among the 200 models generated, the model with lowest objective function was selected for further work. 90,000 compounds from NCI-open database were virtually screened against the switch II pocket of the RanGDP model, using software GOLD (The Cambridge Crystallographic Data Centre, Cambridge, UK).

### Medicinal chemistry

28 top-ranking candidate compounds were kindly provided, as a gift, by the Drug Synthesis and Chemistry Branch, Developmental Therapeutics Program, NCI, Bethesda, MD, USA. Initial hit M26 was re-synthesized and characterized by NMR analysis. Chemical optimization of M26, leading to compound M36, is described in the supplemental information.

### Cell lines and cell culture

The human EOC cell lines used [TOV81D, TOV112D, OV90, OV866(2), TOV1946, OV1946, TOV2295(R), OV4485] were derived in our laboratory from patients’ tumors (TOV) or ascites (OV) ^65–68^. All EOC cell lines were maintained in low oxygen at 7% O_2_ and 5% CO_2_ and grown in OSE medium (Wisent, Montreal, QC) supplemented with 10% FBS (Wisent), 0.5 µg/mL amphotericin B (Wisent) and 50 µg/mL gentamicin (Life Technologies Inc., Burlington, ON). The human retinal epithelial cell line ARPE (ARPE-19) was purchased from American Type Culture Collection (ATCC, Manassas, VA) and maintained in DMEM-F12 (Wisent) medium supplemented with 10% FBS (Wisent), 0.5 µg/mL amphotericin B (Wisent) and 50 µg/mL gentamicin (Life Technologies Inc.).

### Small interference RNA (siRNA) treatment

Suspensions of 10^6^ cells in 100 µL of nucleofector solution V (Lonza Group Ltd, Basel, Switzerland) were transfected by electroporation with 1.2 nmoles siRNA targeting Ran (J-010353-06, ON-TARGETplus, Dharmacon Thermo Fisher Scientific Inc., Waltham, MA). For each experiment, efficiency of Ran silencing was verified 48 hours after transfection by Western blot analysis. Scramble siRNA (D-001810-02, Dharmacon) was used as a control in all experiments.

### Clonogenic survival

Clonogenic assays were performed as previously described ^64^. Colonies were counted under a stereo microscope and reported as percent of control. IC_50_ values were determined using Graph Pad Prism 5 software (GraphPad Software Inc., San Diego, CA). Each experiment was performed in duplicate and repeated three times. Sensitivity of the cell lines to small molecule inhibitors of Ran was assessed using a concentration range of 0–40 µM.

### IncuCyte cell proliferation phase-contrast imaging assay

Cells (2,000 cells/well) were plated in a 96-well plate. The next day, compounds and/or Olaparib (S1060, Cedarlane) were added at the indicated concentrations. Following treatment, cell confluence was imaged by phase contrast using the IncuCyte live cell monitoring system (Essen BioScience, Ann Arbor, MI). Frames were captured at day 0 (immediately after treatment) and at the end of the experiment using a 10X objective. For Ran KD experiments, cells were seeded in a 96-well plate (4,000 cells/well) directly after transfection. Cell confluence monitoring started the next day as described above.

### Protein preparation and western blot analysis

Cells were lysed with RIPA buffer containing protease inhibitors. Whole cell lysates were run through a Bradford assay (Thermo Fisher Scientific) for protein quantification. Around 25–50 µg of protein was separated by 12.5% SDS-PAGE and transferred onto nitrocellulose membranes. The resultant blots were probed with Ran (1:10 000, sc-271376 Santa Cruz Biotechnology, Dallas, TX), cleaved PARP (1:1000, #9541, Cell Signaling Technology Inc., Danvers, MA), cleaved caspase 3 (1:1000, #9661, Cell Signaling Technology Inc.), GAPDH (1:2500, #2118, Cell Signaling Technology Inc.) or beta-actin (1:50 000, ab6276, Abcam Inc., Toronto, ON, Canada) primary antibodies overnight at 4°C then with peroxidase-conjugated secondary antibodies for 2 hours at room temperature. Proteins were detected using enhanced chemiluminescence (Thermo Fisher Scientific).

### RT-PCR

The total RNA from TOV-112D or OV1946 cells was isolated using RNeasy Kit (Qiagen®). The total RNA concentration and purity were measured on a NanoDrop™ spectrophotometer. RNA was reverse transcribed using the QuantiTect Reverse Transcription Kit (Qiagen®) according to the manufacturer’s protocol. cDNA amplification was performed with SYBR Green PCR master mix (Applied Biosystems®) using the StepOnePlus Real-Time PCR system (Applied Biosystem®). Negative controls were included in all experiments, and actin served as the housekeeping gene. Primers were ordered from Integrated DNA Technologies, Inc:

NR1D1:

Forward primer: TGGACTCCAACAACAACACAG
Reverse primer: GATGGTGGGAAGTAGGTGGG

β-actin:

Forward primer: CATGTACGTTGCTATCCAGGC
Reverse primer: CTCCTTAATGTCACGCACGAT

### Cell death analysis by flow cytometry

Cells were treated with M36 and/or Olaparib. Six days after treatment, adherent and floating cells were mixed and incubated 5 minutes at room temperature with DRAQ 7 (ab109202, Abcam Inc). A maximum of 30,000 events were counted per condition using the Fortessa flow cytometer (BD Biosciences, Mississauga, ON) and analyzed with the FlowJo software.

### Immunofluorescence

M36 and/or Olaparib-treated and control cells grown on coverslips were washed with 1X PBS, fixed in 4% paraformaldehyde and permeabilized with 0.25% Triton X-100 (Sigma-Aldrich Inc.). After blocking (4% BSA and 4% FBS in PBS), coverslips were incubated with anti– RanGTP (26915, NewEastBioscences, diluted 1:100), anti p-γ-H2AX (Ser139) (05-636, EMD Millipore, diluted 1:2000), anti-Rad51 (ab213, Abcam, diluted 1:750), anti 53BP1 (NB100-305, Novus Biologicals, diluted 1:2500) and anti-Geminin (10802-1-AP, Proteintech, diluted 1:1000) antibodies in blocking buffer for 2 hours at room temperature. Subsequently, samples were incubated with Cy-5 (1:500, Life Technologies Inc.) and Alexa Fluor 488 (for Geminin) secondary antibodies for 1 hour. Coverslips were mounted onto slides using Prolong® Gold anti-fade reagent with DAPI (Life Technologies Inc.). Images were obtained using a Zeiss microscope (Zeiss observer Z1) with a 20X objective. For Rad51 and 53BP1 analysis, cells were transfected with siRan or treated with M36 for 48 hours then gamma-irradiated at 10 Gy and fixed one hour after irradiation. Automated analysis software from Zeiss (AxioVision™, Carl Zeiss) was used for foci counting. In each condition, γ-H2AX, Rad51 and 53BP1 foci were quantified in roughly 400 nuclei.

### Cellular Thermal Shift Assay (CETSA)

OV1946 cells (10^7^ cells/petri) were seeded in 100 mm × 15 mm petri dish and allowed to attach for 24 h. Then, cells were treated with DMSO or M36 at concentrations from 50 to 200 μM and incubated for 1 h at 37 °C. After incubation, cells were washed, trypsinized and centrifuged for 5 minutes at 1200 rpm. Cells were resuspended in HBSS, divided into different aliquots and heated for three minutes at different temperatures using the Biometra TOne PCR (Analytikjena, Jena, Germany). Cell lysates were then obtained by three freeze-thaw cycles in liquid nitrogen (1 min each step of each cycle) followed by centrifugation at 20,000 × g for 20 min at 4°C. The supernatants were harvested, and SDS-PAGE loading buffer was added before boiling for 5min. Proteins were migrated and blotted into PVDF membranes. Primary antibodies used were: Ran (ab53775, Abcam, 1/5000) and β-Actin (ab6276, Abcam, 1/5000). Proteins were detected by HRP-secondary antibody and the ECL detection kit (Clarity and Clarity Max ECL Western Blotting Substrates from Biorad). Densitometry analyses were conducted using the UV-VIS Spectrophotometer (Thermo Fisher Scientific) and relative intensity of Ran protein was obtained by the ratio over the β-actin band.

### HiBiT CETSA

The cDNA of human Ran was cloned into the mammalian expression vector pcDNA3.1(+) with an N-terminal HiBiT tag (from Promega) for luminescence detection. The empty pcDNA3.1(+) vector was used as negative control. OV1946 cells (4 × 10^6^ cells/well) cells were seeded in 6-well plates. After 24 h, transfection was performed using Lipofectamine™ 3000 (Thermo Fisher Scientific) according to the manufacturer’s protocol. Twenty-four hours post-transfection, cells were treated with DMSO (vehicle control) or 100 µM M36 for 1 h. Cells were then harvested, resuspended in HBSS, and distributed into 96-well PCR plates. Samples were heated at different temperatures for 3 min in a thermocycler and rapidly cooled on ice. Following heat treatment, cells were lysed using the Nano-Glo® HiBiT Lytic Detection System (Promega) and incubated for 10 min at room temperature. Luminescence was measured according to the manufacturer’s instructions. Luminescence readings were normalized to the background value of the empty plasmid and to the value of the first temperature used.

### Ran, Cdc42, RhoA and Rac1 activation assays

Cells were seeded onto 6-well tissue culture plates in such a way that cell confluence reached approximately 70% the day of the experiment. The day of the experiment cells were treated for 1 hour with the indicated compounds prior to protein extraction and quantification. Assays were performed using the Ran, Rac1 activation assay kits (Cell Biolabs) and the RhoA and Cdc42 G-LISA activation assay kits (Cytoskeleton®) according to the manufacturer recommendations.

#### In vivo study

For the PK studies, 6-week-old female CD1 mice (Charles River laboratories, Senneville, QC, Canada) received a single intraperitoneal injection of M36 (200 mg/kg), dissolved in DMSO 10%, Kolliphor® EL 10%, PEG-400 20% and PBS 60%. For each time point (5, 30, 120, 240, 360 and 1440 minutes), 3 mice were sacrificed and blood was collected by cardiac puncture. Plasma levels of the compound were measured by mass spectrometry. For the tolerance test, M36 was dissolved as described above and injected intraperitoneally into 6-week-old female NOD.Cg-Rag1^tm1Mom^Il2rg^tm1Wjl^/SzJ (NRG) female mice (The Jackson laboratory, Bar Harbor, ME) every day (except on weekends) at 200 mg/kg. During this study, mice were monitored for survival and weight loss/gain. For the tumor xenograft experiment, NRG mice were injected with 10^6^ TOV112D cells suspended in a mix of 1:1 PBS and matrigel (BD Biosciences) at subcutaneous sites. Treatments with M36 or Olaparib started the following day. Measurements of tumor size were collected at least twice a week until tumors reached limit points (2000 mm3). All animal procedures were performed in accordance with the guidelines for the Care and Use of Laboratory Animals of the CRCHUM and were approved by the Comité institutionnel de protection des animaux (CIPA) (Protocols #C18009AMMs and C22014AMMs).

### Micro-dissected tissue (MDT) production, culture and treatment

MDTs were obtained from both EOC cell xenografts or patient tissue samples. For the xenografts, cell suspensions containing 1×10^6^ cells for TOV112D or 5×10^6^ cells for OV1946 in PBS-Matrigel were injected subcutaneously in NRG female mice. Solid tumors were harvested at volumes between 800 and 1500 mm^3^. For patient tissue, tumor sample was collected following informed consent from the Centre hospitalier de l’Université de Montréal (CHUM), Division of Gynecologic Oncology. Tissue was processed immediately upon retrieval. The use of human sample was approved by the Comité d’éthique de la recherche du CHUM (CÉR-CHUM, Protocol #BD 04.002).

The micro-dissection and loading procedures were adapted from previously published work ^46,69^. Briefly, both xenograft and patient tumor sample were sliced into 1cm slices using a scalpel and processed into 350µm slices using a McIlwain™ tissue chopper (Ted Pella©). Tissue slices were kept in Hank’s balanced salt solution (HBSS, Wisent) supplemented with 10% FBS, 55 µg/mL of gentamicin and 0.6 µg/mL of amphotericin B. A 500 µm biopsy punch (PUN0500, Zivic instruments) was then used to produce MDTs, spheroidal tissue of approximately 300 µm in diameter. The loading and trapping of MDTs in microfluidic devices were performed as previously described ^46^. MDTs were cultured in OSE (xenografts) or Ovarian TumorMACS™ (MILTENYI BIOTEC; patient sample) media supplemented with 10% FBS, 55 ug/mL of gentamicin and 0.6 ug/mL of amphotericin B.

MDTs were treated with either 100 or 200 µM of M36 in 1% DMSO (Sigma) for 24 or 72hrs continuously. MDTs were then fixed with 10% formalin (Chaptec) and processed following a previously published paraffin-embedding lithography procedure to create micro-dissected tissue microarray (MDTMA) blocks ^46^.

### MDT Immunofluorescence staining and quantification

MDTMA blocks were cut into 4µm sections and put on Matsunami TOMO® hydrophilic adhesion slides (VWR). Slides were stained for KI67 and cleaved-caspase 3 (CC3) to assess proliferation and cell death, respectively, using the *Discovery Ultra* automated stainer (Ventana Medical System inc. (VMSI)). A pan-cytokeratin (CK) antibody was also used to identify the tumor epithelium of the MDTs. Slides were heated at 60°C for 20 min before staining. Antigen retrieval was carried out with Cell Conditioning 1 solution (VMSI) for 90 min. Mouse anti-KI67 (1:500) (9449S, Cell Signaling), rabbit anti-CC3 (1:200) (9661S, Cell Signaling) and guinea pig anti-pan-CK (1:200) (BP5069, Cedarlane) antibodies were dispensed automatically and slides were incubated for 60 minutes at 37°C. Secondary antibodies, including anti-mouse Alexa 488 (1:250) (A-11001, Life Technologies), anti-rabbit Alexa 750 (1:200) (A-21039, Life Technologies) and anti-guinea pig Alexa 647 (1:200) (706-605-148, Cedarlane) were also dispensed automatically and slides were incubated for 60 minutes at 37°C. A mounting medium with DAPI (MJS Biolynx inc) was used to stain the nuclei. Slides were scanned at 20x with either an Olympus BX61 (Olympus) or an Aperio Versa 200 (Leica) microscope. IF filters used for DAPI, KI67, CC3 and tumor epithelium were DAPI, Alexa-488, Cy7 and Cy5, respectively. Stained MDTs were quantified using the VisiomorphDP software (VisioPharm, Denmark, http://visiopharm.com). Percent of CC3 staining or nuclear K-67 staining in the cytokeratin positive area were quantified and plotted as bar graphs.

### Statistical analysis

Data from at least three independent experiments are presented as the means ± SD. Statistical analyses were done using GraphPad Prism version 10 (GraphPad Sofware Inc., http://graphpad.com). Comparisons were carried out using Student’s t-test. Analysis of MDTs was done using Two-way ANOVA followed by Tukey’s test. Comparisons of survival curves were performed using log-rank test. A p-value of less than 0.05 was considered statistically significant.

## ACKNOWLEDGMENTS

We thank Kim Leclerc Désaulniers and Lise Portelance for technical assistances with the animal experiments and biobank specimen procurement, respectively. We acknowledge the CRCHUM animal facility, the ICM Imaging/Live Imaging platform, the Molecular Pathology core facility of the CRCHUM and the CRCHUM Microfluidic platform. A-MM-M and DP are researchers of the CRCHUM/ICM, which receive support from the Fonds de recherche du Québec – Santé (FRQS). ZB was supported by a MITACS fellowship and by the ICM NathalieMoreau Bursary. SR was supported by a studentship from the Canderel Fund of the Institut du cancer de Montréal (ICM).

## FUNDING

This work was supported by the Canadian Institute for Health Research (CIHR MOP142724 and PJT148642), the Oncopole in collaboration with the FRQS, the Cancer Research Society, Génome Québec and IRICoR (#265877), and the ICM (Fonds Défi Spyder and Anne-Marie Chagnon). Cell lines as well as patient tissue sample used in this study were provided by the CRCHUM ovarian tumor bank, which is supported by Ovarian Cancer Canada (OCC) and by the Banque de tissus et de données of the Réseau de recherche sur le cancer (RRCancer) of the FRQS affiliated with the Canadian Tumor Repository Network (CTRNet).

## AUTHOR CONTRIBUTIONS

ZB, XT, EC, JHW and A-MM-M designed the study. ZB, XT, SR, RG and JK-D performed the experiments. ZB, XT, EC, SR, RG and JK-D analyzed the data. ZB and XT wrote the manuscript. EC, JHW, DP, and A-MM-M supervised the study, provided guidance, and edited the manuscript.

## Notes

### Competing Interest Statement

The authors have declared no competing interest.

### Summary of Updates

A sentence was added in the Abstract.

