## Supplemental Information for "Identification of a new inhibitor of Ran GTPase with potential therapeutic value in epithelial ovarian cancer"

**Supplemental Material:**

Supplemental Figure S1

Supplemental Figure S2

Supplemental Figure S3

Supplemental Figure S4

Supplemental Information: Chemical Synthesis and Characterizations

A)

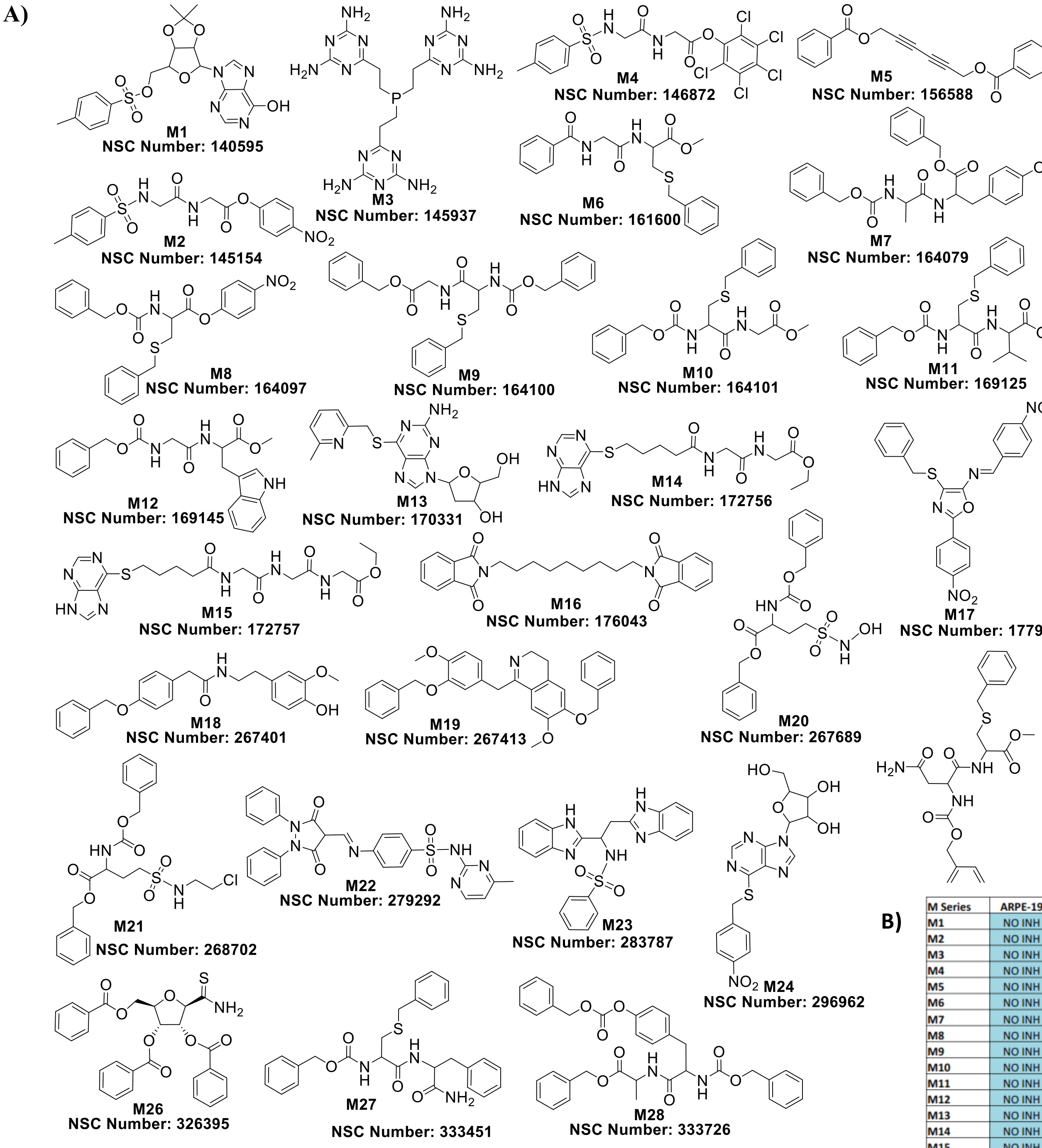


B)

| M Series | ARPE-19 | TOV112D |
|---|---|---|
| M1 | NO INH | NO INH |
| M2 | NO INH | NO INH |
| M3 | NO INH | NO INH |
| M4 | NO INH | NO INH |
| M5 | NO INH | NO INH |
| M6 | NO INH | NO INH |
| M7 | NO INH | NO INH |
| M8 | NO INH | NO INH |
| M9 | NO INH | NO INH |
| M10 | NO INH | NO INH |
| M11 | NO INH | NO INH |
| M12 | NO INH | NO INH |
| M13 | NO INH | NO INH |
| M14 | NO INH | NO INH |
| M15 | NO INH | NO INH |
| M16 | NO INH | NO INH |
| M17 | INH | INH |
| M18 | NO INH | NO INH |
| M19 | NO INH | NO INH |
| M20 | NO INH | NO INH |
| M21 | NO INH | NO INH |
| M22 | NO INH | NO INH |
| M23 | NO INH | NO INH |
| M24 | NO INH | NO INH |
| M25 | NO INH | NO INH |
| M26 | NO INH | INH |
| M27 | NO INH | NO INH |
| M28 | INH | INH |

**Figure S1. (A)** Chemical structures of the 28 top-ranking compounds from the in-silico screening of the NCI open database against the switch II pocket of RanGDP, using software GOLD. **(B)** Cumulative results of colony formation inhibition by these compounds at 10 μM in normal ARPE and EOC TOV112D cells. Only the compound M26 inhibited TOV112D without affecting ARPE (green arrow)

**A**

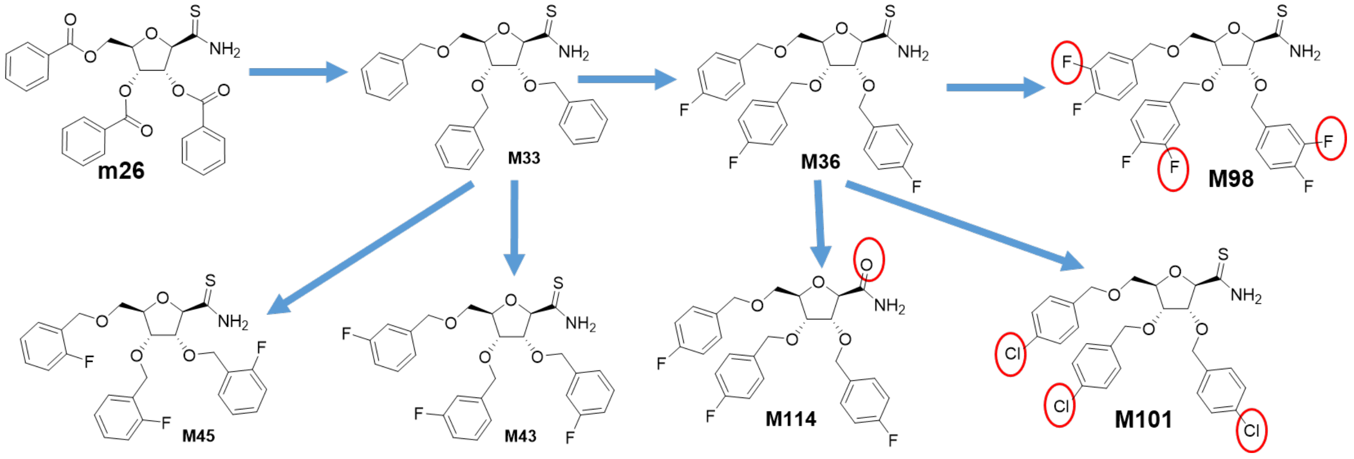


**B**

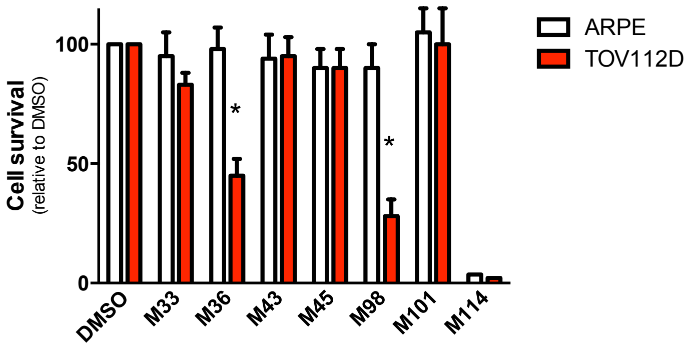


**C**

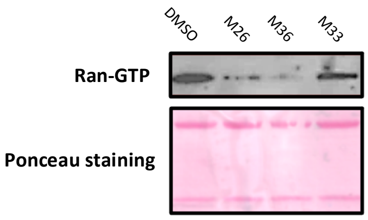


**Figure S2.** Chemical optimization of M26, leading to compound M36. **(A)** Chemical modifications. **(B)** Biological activities of M compounds were tested (10 µM) by clonogenic assay in normal ARPE and TOV112D cells. **(C)** TOV112D cells were incubated for one hour with M compounds (100 µM) before performing RanGTP pull-down assay on the resultant cell extracts.

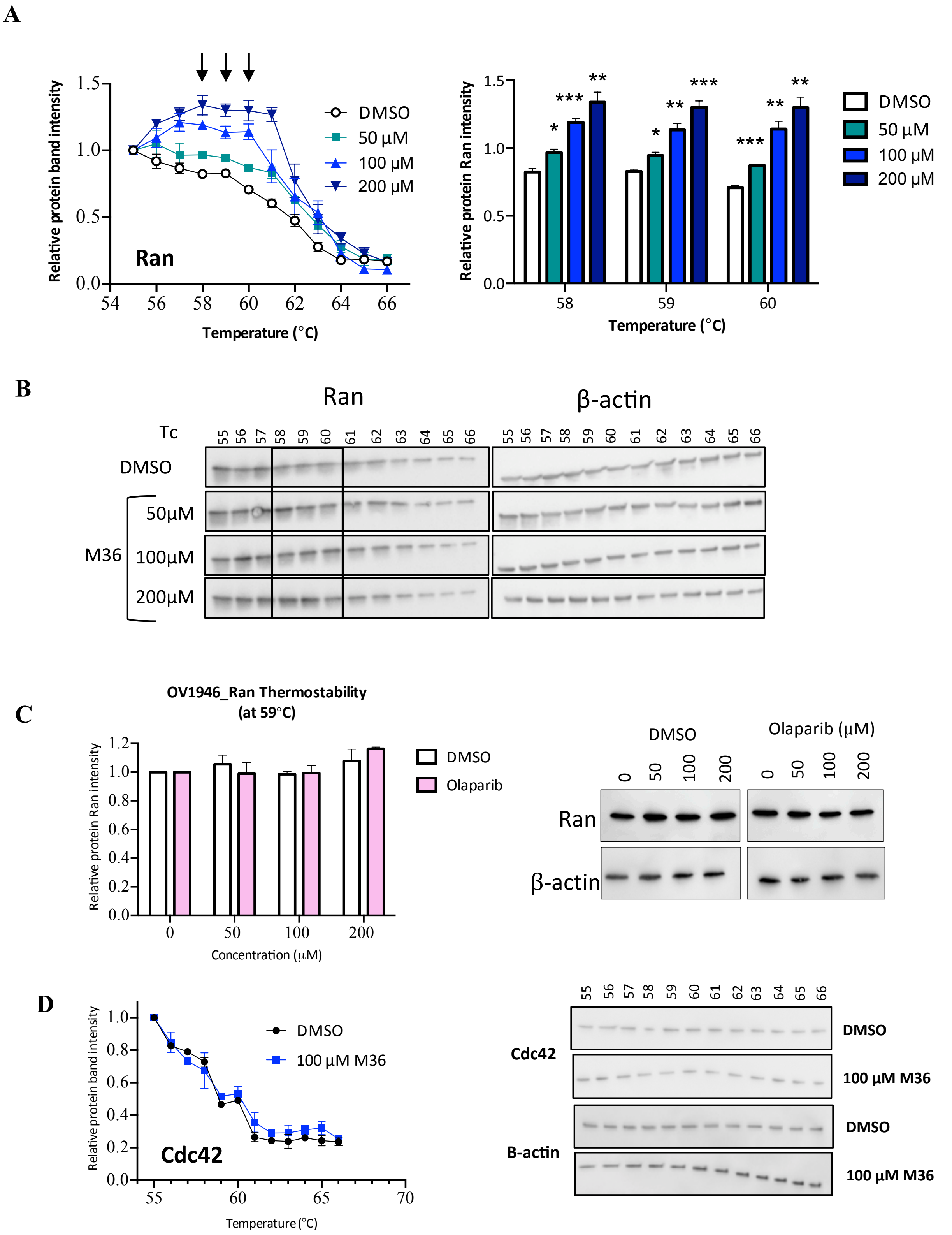

A
Relative protein band intensity
Ran
DMSO
50 μM
100 μM
200 μM
Temperature (°C)
Relative protein Ran intensity
*
***
**
Temperature (°C)
B
Ran
β-actin
Tc
DMSO
50μM
M36
100μM
200μM
C
OV1946_Ran Thermostability
(at 59°C)
Relative protein Ran intensity
DMSO
Olaparib
Concentration (μM)
DMSO
Olaparib (μM)
Ran
β-actin
D
Relative protein band intensity
DMSO
100 μM M36
Cdc42
Temperature (°C)
Cdc42
DMSO
100 μM M36
B-actin
DMSO
100 μM M36

**Figure S3.** Thermo stability evaluation by CETSA. **(A)** OV1946 cells were treated with M36 (50, 100, 200 µM) or DMSO for one hour. CETSA was performed after heating the cell suspensions at different temperatures for three minutes. Cell lysates were subjected to Western blot for evaluation of Ran levels. Data points are densitometry values relative to ß-actin control and normalized to the levels obtained at the lowest temperature used. Arrows indicate the temperatures where significant values were obtained with all three M36 concentrations when compared to control DMSO (as illustrated in the bar graph). *P < 0.05 **P < 0.01 *** P < 0.001 (n = 3, Student's t-test). **(B)** Representative images of the Western Blots; black rectangular box encompasses the results obtained with the three above-mentioned temperatures. **(C)** OV1946 cells were treated for one hour with DMSO or increasing concentrations of Olaparib. CETSA was then performed at 59 °C for three minutes and Ran levels were measured by Western blot. Bar graphs represent densitometry values relative to ß-actin control and normalized to the levels obtained for the untreated cells. Data are from three individual experiments. **(D)** OV1946 cells were treated with 100 µM M36 or DMSO for one hour. CETSA was performed as described above but Cdc42 levels were evaluated by Western blot instead of Ran. Data are from two individual experiments.

A

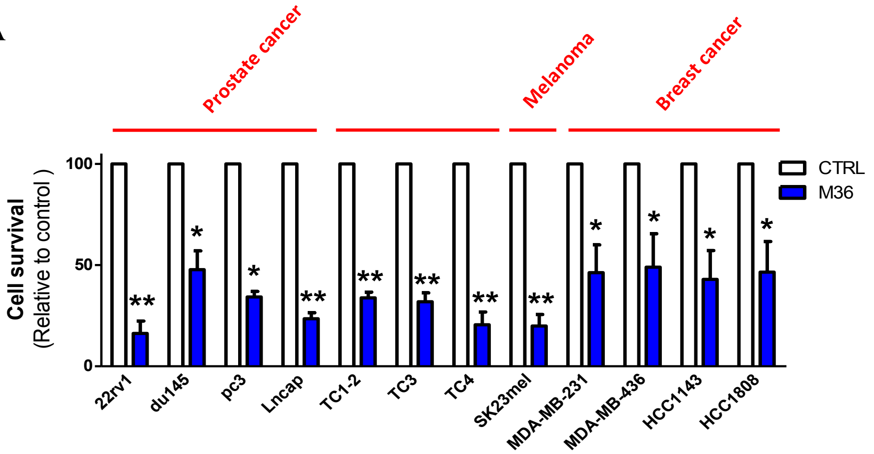
Prostate cancer
Melanoma
Breast cancer
Cell survival
(Relative to control )
100
50
0
CTRL
M36
**
*
*
**
**
**
**
**
*
*
*
*
22rv1
du145
pc3
Lncap
TC1-2
TC3
TC4
SK23mel
MDA-MB-231
MDA-MB-436
HCC1143
HCC1808

B

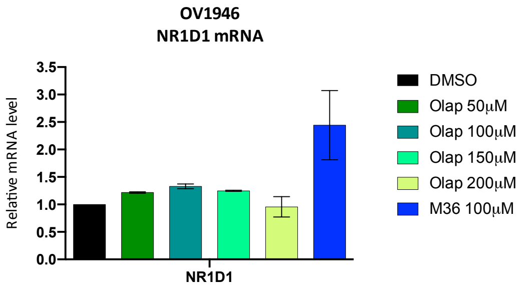
OV1946
NR1D1 mRNA
Relative mRNA level
3.5
3.0
2.5
2.0
1.5
1.0
0.5
0.0
NR1D1
DMSO
Olap 50μM
Olap 100μM
Olap 150μM
Olap 200μM
M36 100μM

C

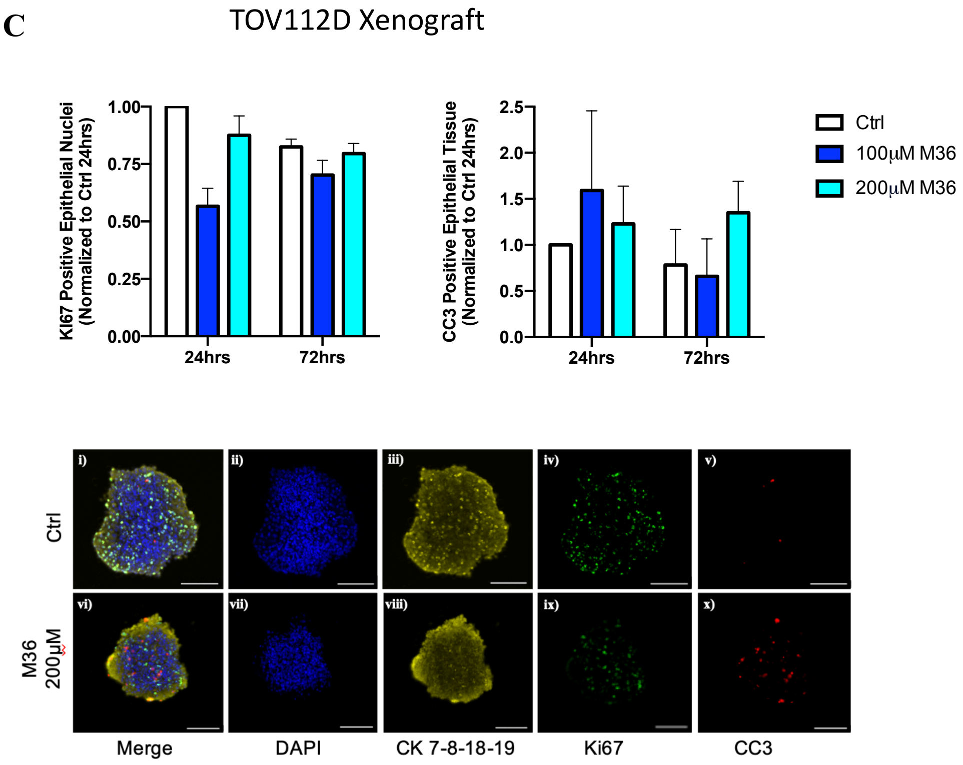
TOV112D Xenograft
Ki67 Positive Epithelial Nuclei
(Normalized to Ctrl 24hrs)
1.00
0.75
0.50
0.25
0.00
24hrs
72hrs
CC3 Positive Epithelial Tissue
(Normalized to Ctrl 24hrs)
2.5
2.0
1.5
1.0
0.5
0.0
24hrs
72hrs
Ctrl
100μM M36
200μM M36
i)
ii)
iii)
iv)
v)
vi)
vii)
viii)
ix)
x)
Ctrl
M36
200μM
Merge
DAPI
CK 7-8-18-19
Ki67
CC3

**Figure S4: (A)** Several cell lines of prostate, gastrointestinal, skin and breast cancers were treated with M36 (40 μM) and subjected to proliferation assays using the IncuCyte live cell monitoring system. Data are expressed as the percentage of DMSO-treated cells at the end of experiment (96 h post-treatment) and are representative of at least three independent experiments. $*P < 0.05$, $**P < 0.01$ ($n \geq 3$, Student's t-test). **(B)** The expression of NR1D1 was evaluated by qRT-PCR in OV1946 cells treated with different concentrations of Olaparib or with 100 μM M36 (positive control) or DMSO (vehicle control). Data are from two individual experiments. **(C)** MDTs from TOV112D xenografts were treated with M36 at different concentrations for 24 or 72 h and then formalin-fixed, paraffin embedded. Sections (6μ) were stained for KI67 (green), cleaved-caspase 3 (red), a cytokeratin cocktail (yellow) and DAPI (blue). Bar graphs represent the expression of nuclear KI67 (left panel) or CC3 (right panel) within the tumour epithelium (CK positive cells) of each MDT. Data shown represents mean ± SEM of two separate experiments, normalized to Ctrl 24hrs. Scale bars of representative images are 200 μm.

### Chemical Synthesis and Characterizations

#### General Experiment Procedure

All reactions were performed in flame-dried glassware under an inert atmosphere of N2. Air- and moisture-sensitive liquids were transferred via syringe. Solutions were concentrated by rotary evaporation at or below 40 °C. Thin-layer chromatography (TLC) was performed on TLC AL foils (silica gel 60 matrix, with fluorescent indicator 254 nm, Sigma-Aldrich, 60778-25EA). TLC plates were visualized by exposure to ultraviolet (UV) light. Flash column chromatography was performed on silica gel (Sigma-Aldrich, 230 – 400 mesh). Solvents and chemical reagents were purchased from Sigma-Aldrich and Asta Tech with > 95% purity which were used without further purification. Proton nuclear magnetic resonance (1H NMR) spectra and carbon nuclear magnetic resonance (13C NMR) spectra were recorded in $CDCl_3$ on a Bruker AvanceIII HD instrument (400 MHz/101 MHz) instrument or Varian Inova 500 instrument (500 MHz/126 MHz) at 23 °C using tetramethylsilane (TMS) as the internal standard. Proton chemical shifts are expressed in parts per million (ppm, $\delta$ scale) and are referenced to residual protium in the NMR solvent ($CHCl_3$: $\delta$ 7.26). Carbon chemical shifts are expressed in parts per million (ppm, $\delta$ scale) and are referenced to the carbon resonance of the NMR solvent ($CDCl_3$: $\delta$ 77.0). Data are repoted as follows: chemical shift, multiplicity (s = singlet, d = doublet, t = triplet, q = quartet, dd = doublet of doublets, dt = doublet of triplets, m = multiplet, br = broad, app = apparent), integration, and coupling constant ($J$) in Hertz (Hz). Accurate (high-resolution) mass spectra were obtained using an Agilent 6224 Accurate-Mass TOF LC/MS mass spectrometer. **M26** was prepared according to literatures[1,2].

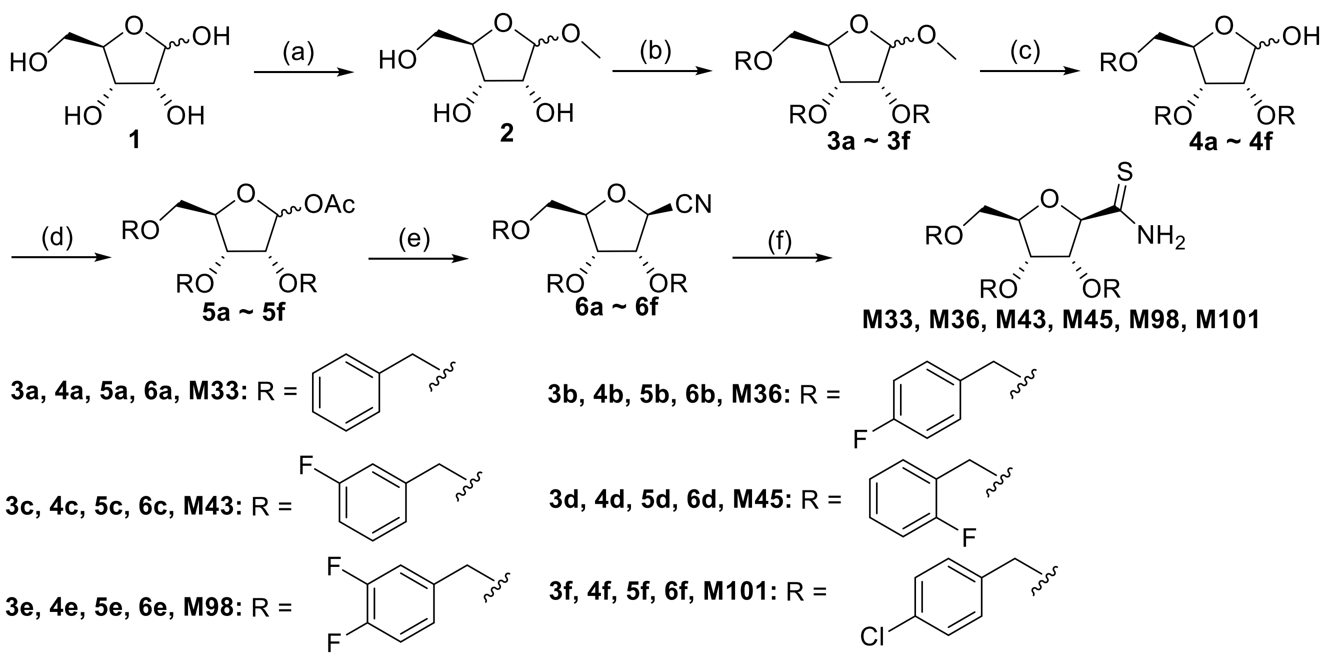

**Scheme S1. Preparation of M33, M36, M43, M45, M98, M101**. (a) AcCl, MeOH, rt, Overnight; (b) substituted benzyl chloride / bromide, Tetrabutylammonium hydrogen sulfate, KOH, THF, reflux,12h; (c) AcOH, 6 mol/L HCl, 65 °C, 1.5h; (d) Pyridine, $Ac_2O$; (e) TMSCN, $BF_3$ $OEt_2$, - 48 °C, 15 min; (f) $H_2S$, DMAP, dry EtOH.

The experimental procedures for the intermediates **3a~3f**, **4a~4f**, **5a~5f**, **6a~6f** have been describe previously[2].

**General procedure for the preparation of M33, M36, M43, M45, M98, M101.**

To a suspension of cyanide **6a~6f** (0.6 mmol) in dry EtOH (20 mL), *N, N*-dimethylaminopyridine (78 mg, 0.06 mmol) was added in one portion under $N_2$. Hydrogen sulfide was slowly passed through the reaction mixture at 0 °C for 2 h. Then the flask was sealed and stirring continued at room temperature for 16 h. The reaction was concentrated and purified by column chromatography.

**2, 5-Anhydro-3, 4, 6-tri-O-benzyl-*β*-D-allonthioamid (M33)**: White solid, yield 92%. $^1$H NMR (500 MHz, $CDCl_3$) $\delta$ 9.10 (br, 1H), 7.56 – 7.44 (m, 2H), 7.38 – 7.26 (m, 9H), 7.25 – 7.20 (m, 2H), 7.17 – 7.13 (m, 2H), 7.09 (br, 1H), 4.96 (s, 1H), 4.91 (d, $J$ = 12.1 Hz, 1H), 4.71 (d, $J$ = 12.1 Hz, 1H), 4.50 – 4.45 (m, 2H), 4.38 (d, $J$ = 10.9 Hz, 1H), 4.35 – 4.34 (m, 1H), 4.30 (d, $J$ = 4.5 Hz, 1H), 4.16 (d, $J$ = 11.8 Hz, 1H), 4.00 – 3.90 (m, 2H), 3.63 (d, $J$ = 10.5 Hz, 1H), 1.29 – 1.23 (m, 1H). M33: $^{13}$C NMR (101 MHz, $CDCl_3$) δ 205.00, 137.42, 137.34, 137.03, 128.73, 128.64, 128.43, 128.40, 128.28, 127.98, 127.96, 127.91, 127.88, 88.50, 81.21, 80.09, 74.58, 73.55, 71.98, 70.93, 67.39. HRMS (ESI) m/z calculated for $C_{27}H_{30}NO_4S$ $[M+H]^+$ 464.1896, found: 464.1899.

**2, 5-Anhydro-3, 4, 6-tri-O-(4-fluoro-benzyl)-*β*-D-allonthioamid (M36)**: White solid, yield 83%.$^1$H NMR (500 MHz, $CDCl_3$) *δ* 9.01 (br, 1H), 7.51 – 7.41 (m, 2H), 7.18 – 7.13 (m, 5H), 7.06 – 6.96 (m, 6H), 4.94 (s, 1H), 4.86 (d, *J* = 12.1 Hz, 1H), 4.67 (d, *J* = 12.1 Hz, 1H), 4.48 (d, *J* = 11.0 Hz, 1H), 4.40 – 4.36 (m, 2H), 4.32 – 4.30 (m, 1H), 4.28 (d, *J* = 4.6 Hz, 1H), 4.13 (d, *J* = 11.6 Hz, 1H), 3.94 – 3.89 (m, 2H), 3.60 (d, *J* = 10.5 Hz, 1H). $^{13}$C NMR (101 MHz, $CDCl_3$) δ 204.82, 163.81, 163.79, 163.72, 161.35, 161.27, 133.10, 133.07, 133.03, 133.00, 132.80, 132.77, 130.57, 130.49, 129.68, 129.60, 129.54, 129.46, 115.73, 115.52, 115.43, 115.38, 115.22, 115.17, 88.29, 81.03, 79.99, 74.51, 72.81, 71.21, 70.11, 67.30. HRMS (ESI) m/z calculated for $C_{27}H_{27}F_3NO_4S$ $[M+H]^+$ 518.1613, found: 518.1617.

**2, 5-Anhydro-3, 4, 6-tri-O-(3-fluoro-benzyl)-*β*-D-allonthioamid (M43)**: White solid, yield 82%. $^1$H NMR (500 MHz, $CDCl_3$) *δ* 8.97 (br, 1H), 7.36 – 7.15 (m, 6H), 7.05 – 6.91 (m, 5H), 6.91 – 6.84 (m, 1H), 4.95 (s, 1H), 4.90 (d, *J* = 12.4 Hz, 1H), 4.70 (d, *J* = 12.4 Hz, 1H), 4.53 – 4.42 (m, 3H), 4.36 – 4.35 (m, 1H), 4.31 (d, *J* = 4.6 Hz, 1H), 4.23 (d, *J* = 12.1 Hz, 1H), 4.03 – 3.92 (m, 2H), 3.67 – 3.65 (m, 1H).M43: $^{13}$C NMR (101 MHz, $CDCl_3$) δ 204.70, 164.13, 164.10, 161.68, 161.65, 139.91, 139.84, 139.55, 139.48, 130.33, 130.25, 130.05, 129.98, 129.90, 123.94, 123.91, 123.19, 123.16, 123.11, 123.08, 115.32, 115.10, 115.02, 114.98, 114.81, 114.77, 114.65, 114.50, 114.44, 114.28, 88.28, 81.53, 79.94, 74.77, 72.79, 72.77, 71.29, 71.28, 70.21, 70.19, 67.47. HRMS (ESI) m/z calculated for $C_{27}H_{27}F_3NO_4S$ $[M+H]^+$ 518.1613, found: 518.1619.

**2, 5-Anhydro-3, 4, 6-tri-O-(2-fluoro-benzyl)-*β*-D-allonthioamid (M45)**: White solid, yield 85%. $^1$H NMR (500 MHz, $CDCl_3$) *δ* 9.06 (br, 1H), 7.56 – 7.52 (m, 1H), 7.36 – 7.20 (m, 6H), 7.16 – 6.97 (m, 5H), 4.98 – 4.96 (m, 2H), 4.73 (d, *J* = 12.2 Hz, 1H), 4.60 (d, *J* = 11.4 Hz, 1H), 4.56 – 4.48 (m, 2H), 4.43 (d, *J* = 11.8 Hz, 1H), 4.37 (d, *J* = 4.5 Hz, 1H), 4.33 – 4.31 (m, 1H), 4.04 – 4.02 (m, 1H), 3.98 – 3.96 (m, 1H), 3.66 – 3.63 (m, 1H).**M45:** $^{13}$C NMR (101 MHz, $CDCl_3$) δ 204.73, 162.24, 162.18, 161.94, 159.78, 159.72, 159.48, 130.83, 130.79, 130.41, 130.36, 130.28, 130.25, 130.21, 129.78, 129.74, 129.70, 129.66, 124.70, 124.64, 124.55, 124.50, 124.32, 124.28, 124.24, 124.10, 124.06, 123.99, 123.96, 115.75, 115.54, 115.38, 115.35, 115.17, 115.14, 88.36, 82.31, 79.98, 75.37, 67.40, 67.36, 65.96, 65.92, 65.34, 65.31, 53.47. HRMS (ESI) m/z calculated for $C_{27}H_{27}F_3NO_4S$ $[M+H]^+$ 518.1613, found: 518.1615.

**2, 5-Anhydro-3, 4, 6-tri-O-(3, 4-difluoro-benzyl)-*β*-D-allonthioamid (M98):** White solid, yield 79%. $^1$H NMR (400 MHz, $CDCl_3$) *δ* 8.89 (br, 1H), 7.39 – 7.27 (m, 2H), 7.23 – 6.85 (m, 8H), 4.93 (s, 1H), 4.85 (d, *J* = 12.3 Hz, 1H), 4.64 (d, *J* = 12.3 Hz, 1H), 4.51 (d, *J* = 11.6 Hz, 1H), 4.44 – 4.39 (m, 2H), 4.35 – 4.30 (m, 2H), 4.22 (d, *J* = 11.9 Hz, 1H), 4.01 – 3.91 (m, 2H), 3.65 – 3.62 (m, 1H). $^{13}$C NMR (101 MHz, $CDCl_3$) δ 204.35, 151.70, 151.61, 151.32, 151.18, 149.22, 149.09, 148.97, 148.83, 134.32, 134.28, 134.23, 133.99, 124.47, 124.44, 124.41, 124.37, 123.71, 123.67, 123.64, 123.61, 123.44, 123.40, 123.38, 123.34, 117.69, 117.52, 117.41, 117.34, 117.28, 117.24, 117.17, 117.11, 116.67, 116.49, 116.31, 88.23, 81.45, 79.89, 74.85, 72.32, 70.80, 69.70, 67.45. HRMS (ESI) m/z calculated for $C_{27}H_{24}F_6NO_4S$ $[M+H]^+$ 572.1330, found: 572.1335.

**2, 5-Anhydro-3, 4, 6-tri-O-(4-chloro-benzyl)-*β*-D-allonthioamid (M101):** Colorless syrup, yield 79%. $^1$H NMR (400 MHz, $CDCl_3$) *δ* 8.97 (br, 1H), 7.42 – 7.39 (m, 2H), 7.34 – 7.27 (m, 6H), 7.19 (br, 1H), 7.17 – 7.07 (m, 4H), 4.93 (s, 1H), 4.86 (d, *J* = 12.3 Hz, 1H), 4.66 (d, *J* = 12.3 Hz, 1H), 4.48 (d, *J* = 11.2 Hz, 1H), 4.42 – 4.35 (m, 2H), 4.34 – 4.26 (m, 2H), 4.15 (d, *J* = 11.9 Hz, 1H), 3.97 – 3.87 (m, 2H), 3.60 (d, *J* = 9.8 Hz, 1H). $^{13}$C NMR (101 MHz, $CDCl_3$) δ 204.68, 135.75, 135.73, 135.39, 134.23, 133.87, 133.86, 129.94, 129.15, 129.04, 128.91, 128.63, 128.59, 88.23, 81.26, 79.94, 74.60, 72.76, 71.19, 70.11, 67.37. HRMS (ESI) m/z calculated for $C_{27}H_{27}F_3NO_4S$ $[M+H]^+$ 566.0726, found: 566.0730.

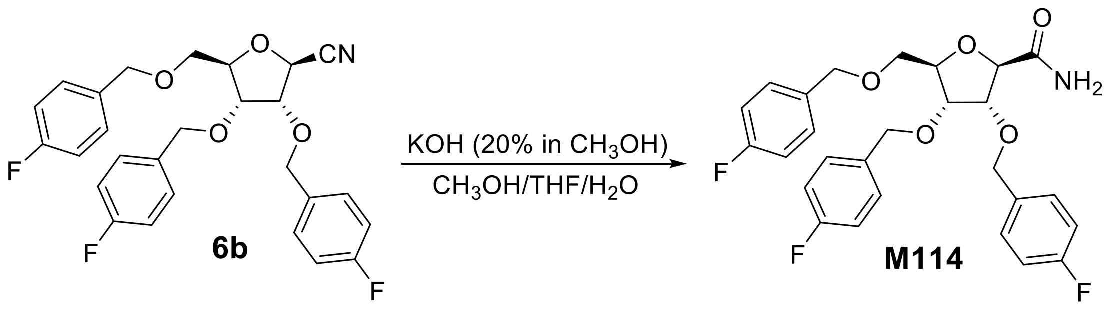

**Scheme S2. Preparation of M114**.

**2, 5-Anhydro-3, 4, 6-tri-O-(4-fluoro-benzyl)-*β*-D-allonamide (M114):** To a solution 2, 3, 5-tri-O-(4-fluoro-benzyl)-*β*-D-ribofuranosylcyanid **6b** (0.24 g, 0.5 mmol) in the mixture of $CH_3OH$ (10 mL), THF (2 mL) and $H_2O$ (2 mL), 3.6 mL of 20 % KOH in $CH_3OH$ was added at room temperature under $N_2$. The reaction mixture was stirred at reflux for 4 h. The reaction mixture quenched by pouring the mixture into ice water and extracting the mixture with $CH_2Cl_2$ (3x). The combined organic layers were washed with brine, dried over anhydrous $Na_2SO_4$, filtered and concentrated under reduced pressure. The residue was purified by flash chromatography (25 – 50 % Ethyl acetate/Hexane) yielding 2, 5-anhydro-3, 4, 6-tri-O-(4-fluoro-benzyl)-*β*-D-allonamide (**M114**) (78 mg, 31 % yield) as a colorless syrup. $^1$H NMR (400 MHz, $CDCl_3$) $\delta$ 7.44 – 7.33 (m, 3H), 7.24 – 7.12 (m, 4H), 7.08 – 6.89 (m, 6H), 5.27 (s, 1H), 4.78 (d, $J$ = 11.9 Hz, 1H), 4.61 – 4.45 (m, 4H), 4.40 (d, $J$ = 11.3 Hz, 1H), 4.28 – 4.23 (m, 2H), 4.14 – 4.13 (m, 1H), 4.01 – 3.97 (m, 1H), 3.89 – 3.85 (m, 1H), 3.59 – 3.56 (m, 1H). $^{13}$C NMR (101 MHz, $CDCl_3$) δ 190.52, 173.78, 163.70, 163.68, 161.26, 161.25, 161.23, 133.25, 133.21, 133.18, 133.16, 133.15, 133.13, 130.12, 130.04, 129.93, 129.74, 129.66, 129.59, 129.54, 129.51, 129.46, 115.55, 115.41, 115.39, 115.34, 115.20, 115.17, 82.28, 79.81, 79.19, 75.62, 72.65, 71.39, 70.58, 67.31, 53.49. HRMS (ESI) m/z calculated for $C_{27}H_{27}F_3NO_5$ $[M+H]^+$ 502.1841, found: 502.1845.


1. Ramasamy, K. S., Averett, D. (1999). A Modified Synthesis of Tiazofurin. Nucleosides and Nucleotides, 18(11–12), 2425–2431.
2. Wu J. H., Batist G., Tian X., Li X., Provencher D, Mes-Masson AM, Carmona E.. Compounds, pharmaceutical compositions and use thereof as inhibitors of ran gtpase. WO2019046931A1

### NMR spectra

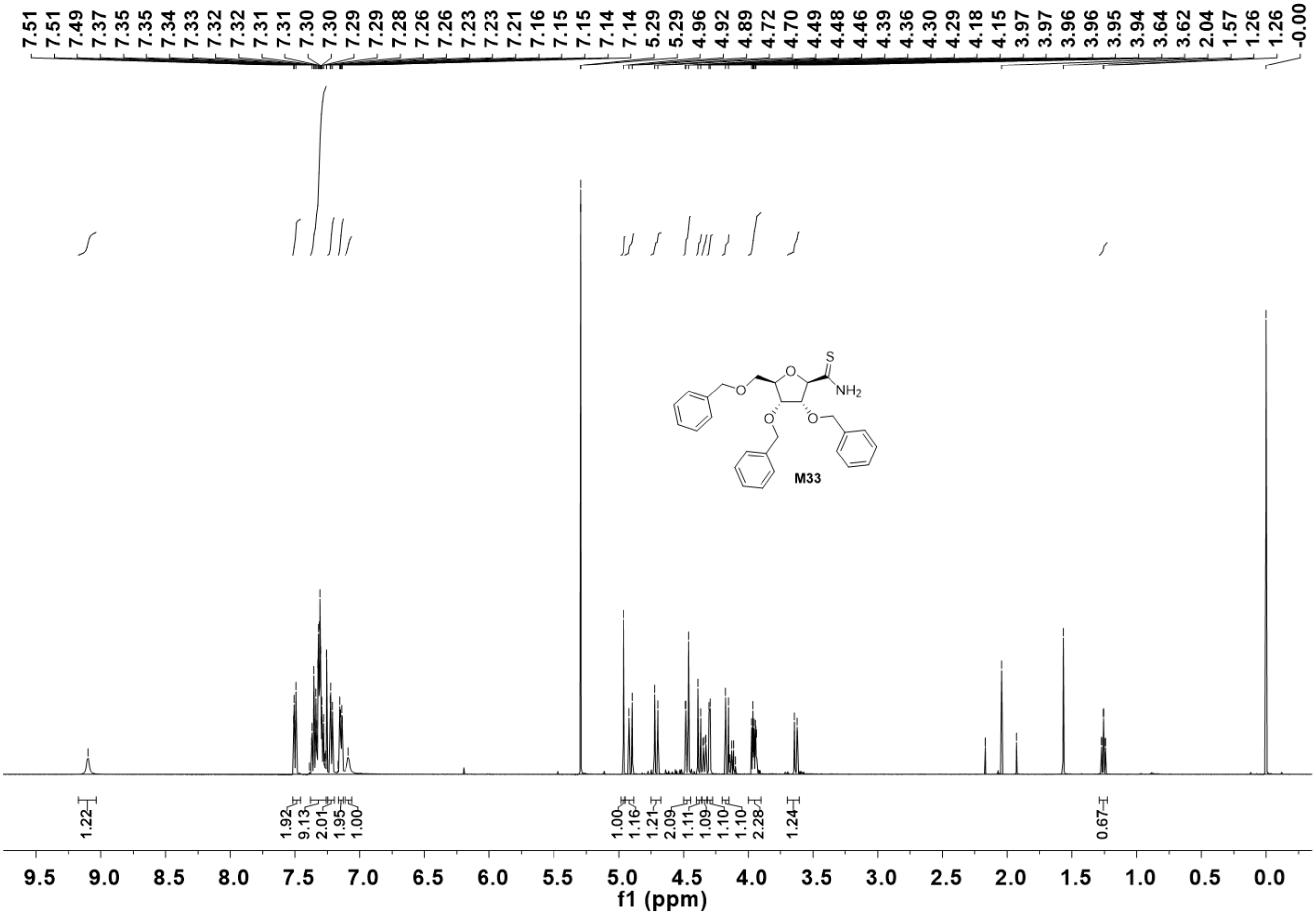
7.51 7.51 7.49 7.37 7.35 7.35 7.34 7.33 7.32 7.32 7.31 7.31 7.30 7.30 7.29 7.29 7.28 7.26 7.26 7.23 7.23 7.21 7.16 7.15 7.15 7.14 7.14 5.29 5.29 4.96 4.92 4.89 4.72 4.70 4.49 4.48 4.46 4.39 4.36 4.30 4.29 4.18 4.15 3.97 3.97 3.96 3.96 3.95 3.94 3.64 3.62 2.04 1.57 1.26 -0.00
S
NH2
O
M33
1.22
1.92 9.13 2.01 1.95 1.00
1.00 1.16 1.21 2.09 1.11 1.09 1.10 2.28
1.24
0.67
9.5 9.0 8.5 8.0 7.5 7.0 6.5 6.0 5.5 5.0 4.5 4.0 3.5 3.0 2.5 2.0 1.5 1.0 0.5 0.0
f1 (ppm)

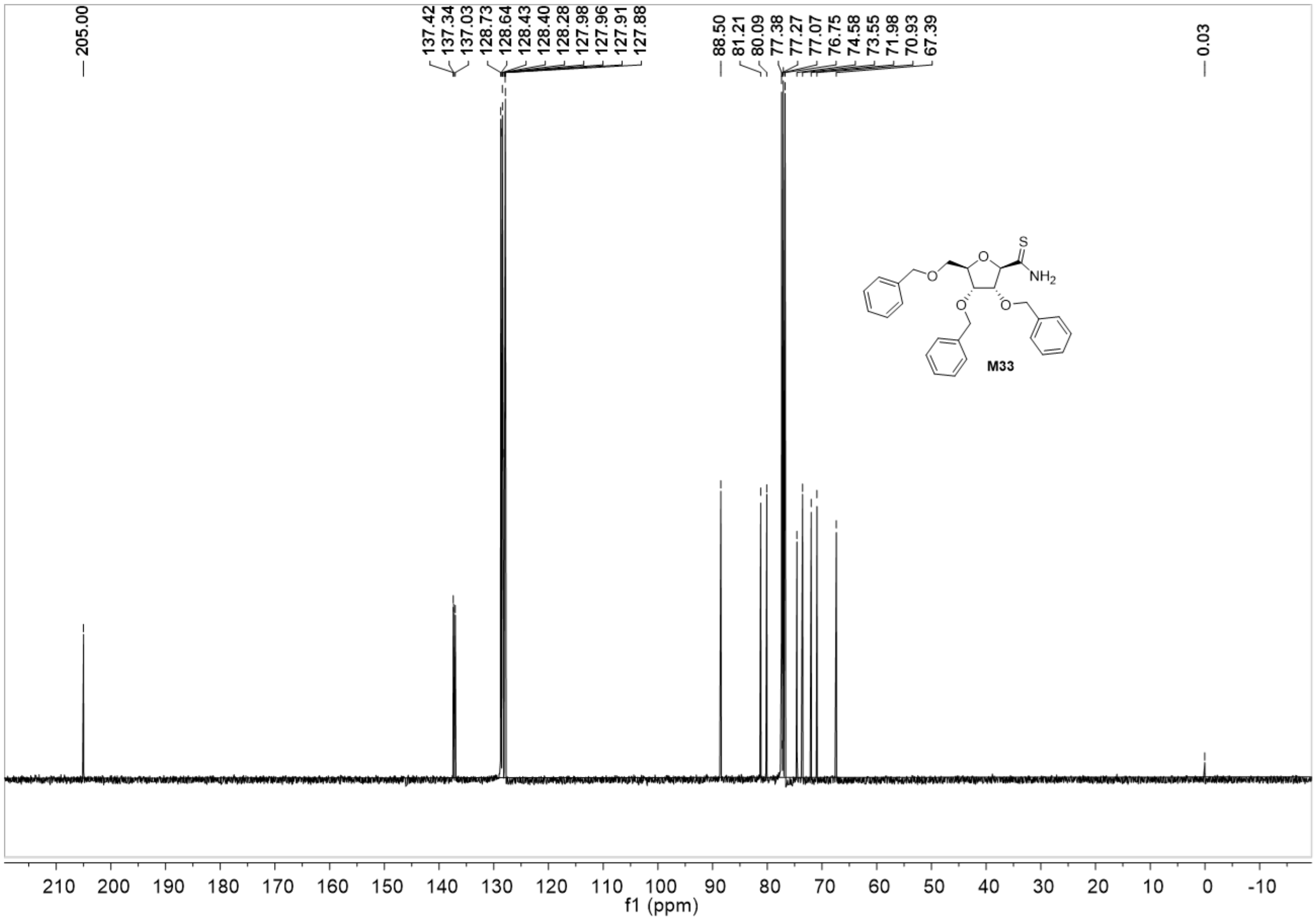
205.00
137.42
137.34
137.03
128.73
128.64
128.43
128.40
128.28
127.98
127.96
127.91
127.88
88.50
81.21
80.09
77.38
77.27
77.07
76.75
74.58
73.55
71.98
70.93
67.39
0.03
S
O
$NH_2$
M33
210 200 190 180 170 160 150 140 130 120 110 100 90 80 70 60 50 40 30 20 10 0 -10
f1 (ppm)

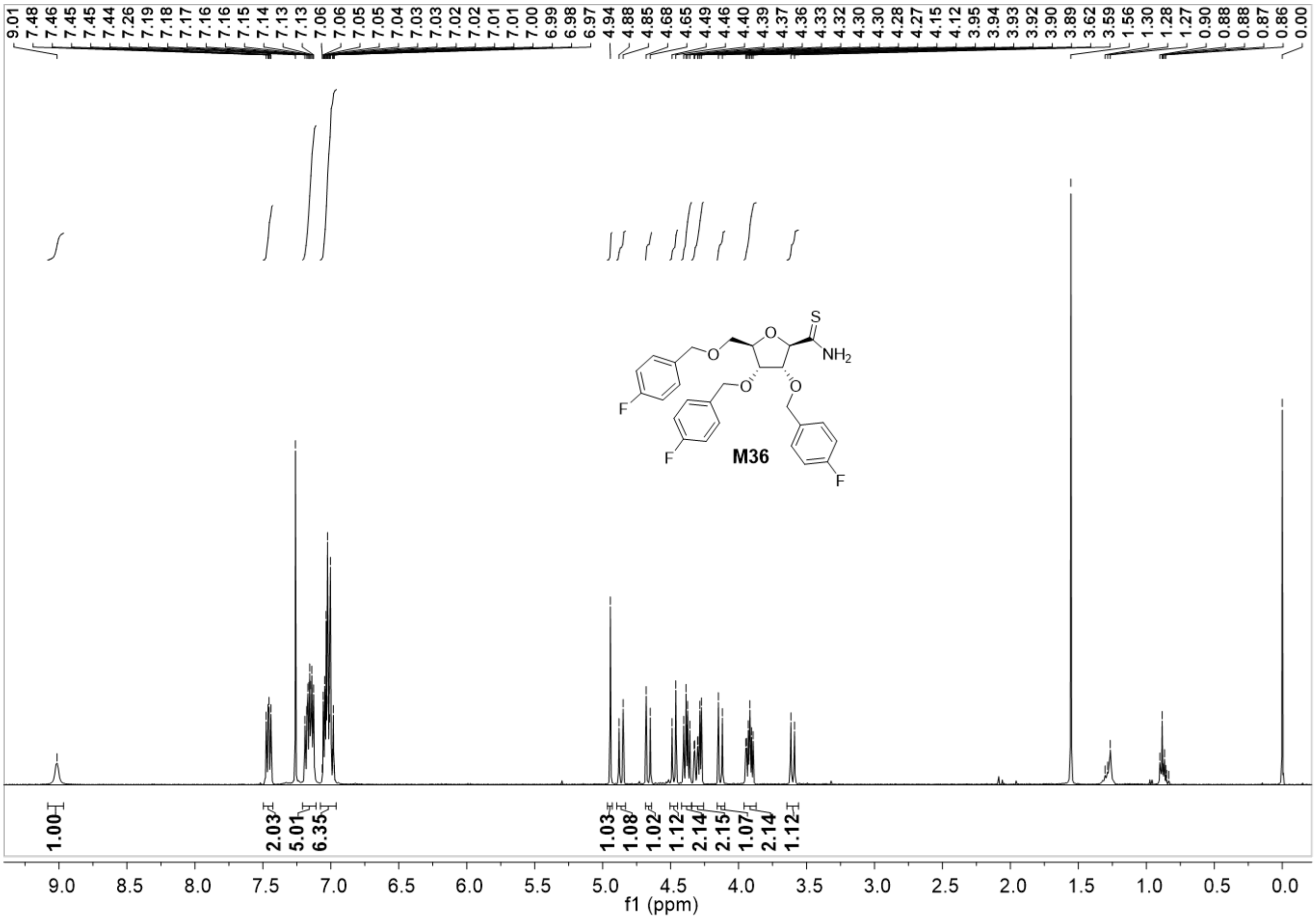
9.01 7.48 7.46 7.45 7.44 7.26 7.19 7.18 7.17 7.16 7.15 7.14 7.13 7.06 7.05 7.05 7.04 7.03 7.02 7.01 7.00 6.99 6.98 6.97 4.94 4.88 4.85 4.68 4.65 4.49 4.46 4.40 4.39 4.37 4.36 4.33 4.32 4.30 4.30 4.28 4.27 4.15 4.12 3.95 3.94 3.93 3.92 3.90 3.89 3.62 3.59 1.56 1.30 1.28 1.27 0.90 0.88 0.88 0.87 0.86 0.00
S
NH2
O
F
F
F
M36
1.00
2.03
5.01
6.35
1.03
1.08
1.02
1.12
2.14
2.15
1.07
2.14
1.12
9.0 8.5 8.0 7.5 7.0 6.5 6.0 5.5 5.0 4.5 4.0 3.5 3.0 2.5 2.0 1.5 1.0 0.5 0.0
f1 (ppm)

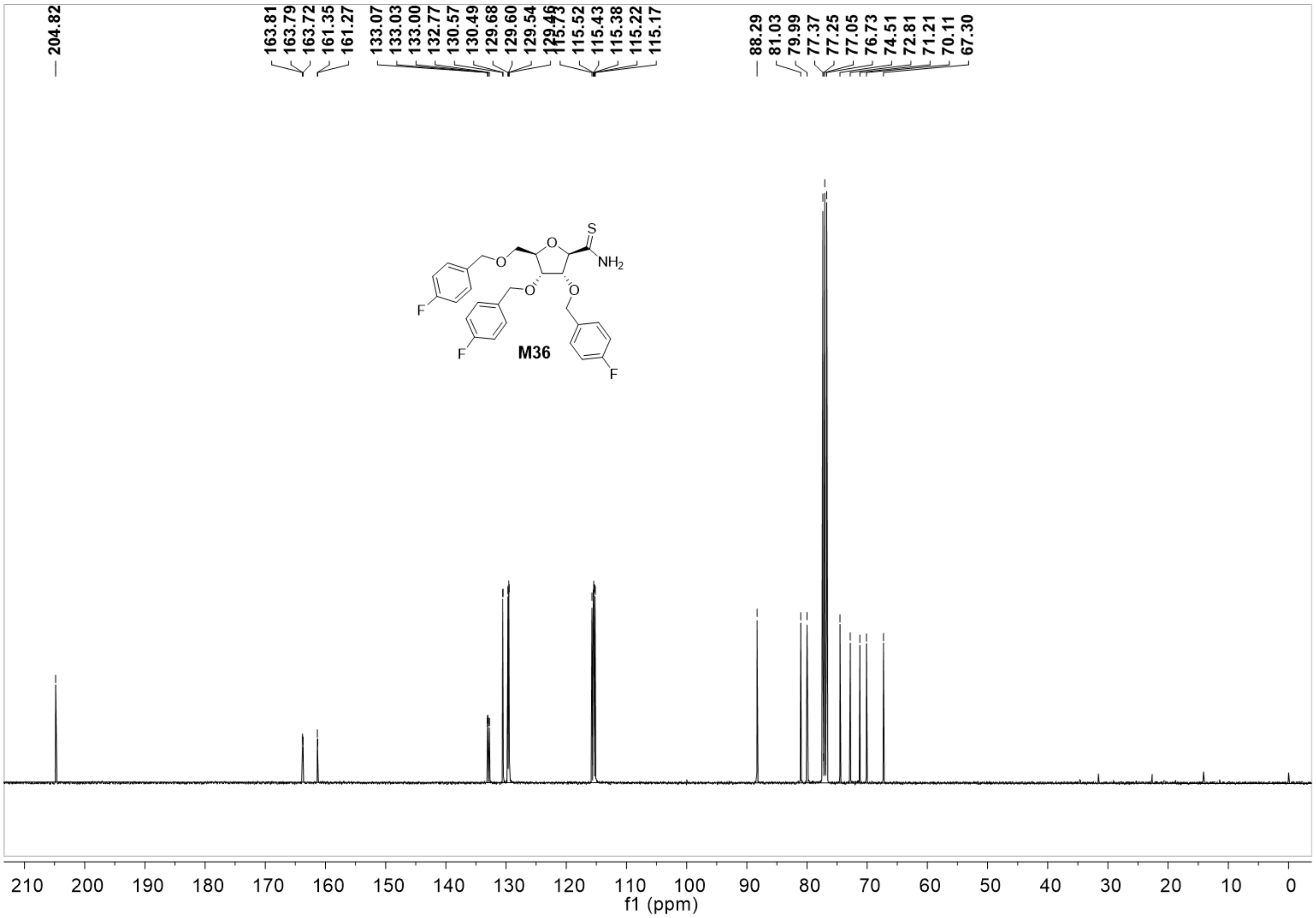
204.82
163.81
163.79
163.72
161.35
161.27
133.07
133.03
133.00
132.77
130.57
130.49
129.68
129.60
129.54
129.46
115.73
115.52
115.43
115.38
115.22
115.17
88.29
81.03
79.99
77.37
77.25
77.05
76.73
74.51
72.81
71.21
70.11
67.30
S
NH2
O
F
M36
210 200 190 180 170 160 150 140 130 120 110 100 90 80 70 60 50 40 30 20 10 0
f1 (ppm)

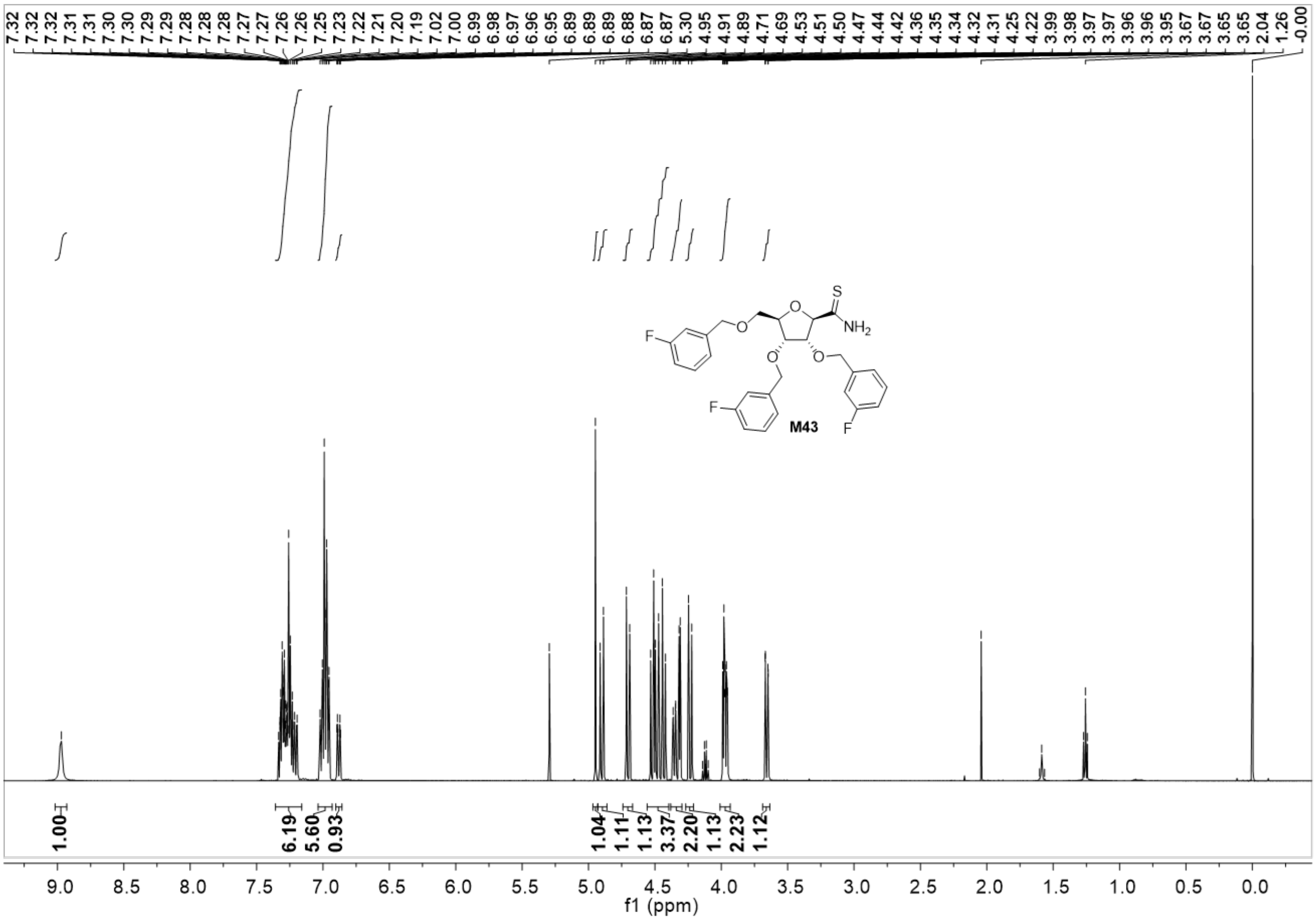

7.32 7.32 7.32 7.31 7.30 7.30 7.29 7.29 7.28 7.28 7.28 7.27 7.27 7.26 7.26 7.25 7.23 7.22 7.21 7.20 7.19 7.02 7.00 6.99 6.98 6.97 6.96 6.95 6.89 6.89 6.88 6.87 6.87 5.30 4.95 4.91 4.89 4.71 4.69 4.53 4.51 4.50 4.47 4.44 4.42 4.36 4.35 4.34 4.32 4.31 4.25 4.22 3.99 3.98 3.97 3.97 3.96 3.96 3.95 3.67 3.67 3.65 3.65 2.04 1.26 -0.00
M43
1.00 6.19 5.60 0.93 1.04 1.11 1.13 3.37 2.20 1.13 2.23 1.12
f1 (ppm)

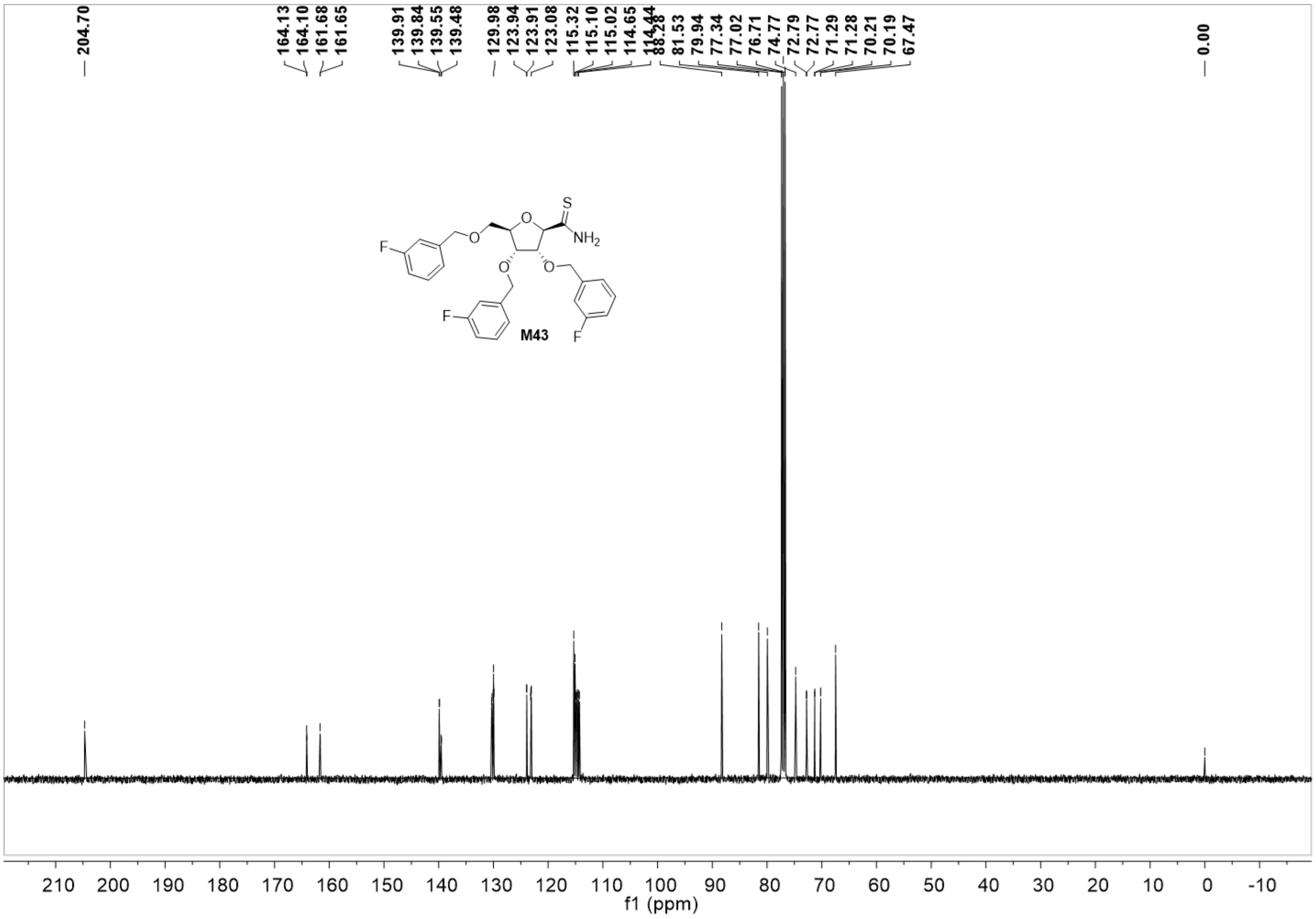

204.70
164.13
164.10
161.68
161.65
139.91
139.84
139.55
139.48
129.98
123.94
123.91
123.08
115.32
115.10
115.02
114.65
114.44
88.28
81.53
79.94
77.34
77.02
76.71
74.77
72.79
72.77
71.29
71.28
70.21
70.19
67.47
0.00
S
O
NH2
F
M43
210 200 190 180 170 160 150 140 130 120 110 100 90 80 70 60 50 40 30 20 10 0 -10
f1 (ppm)

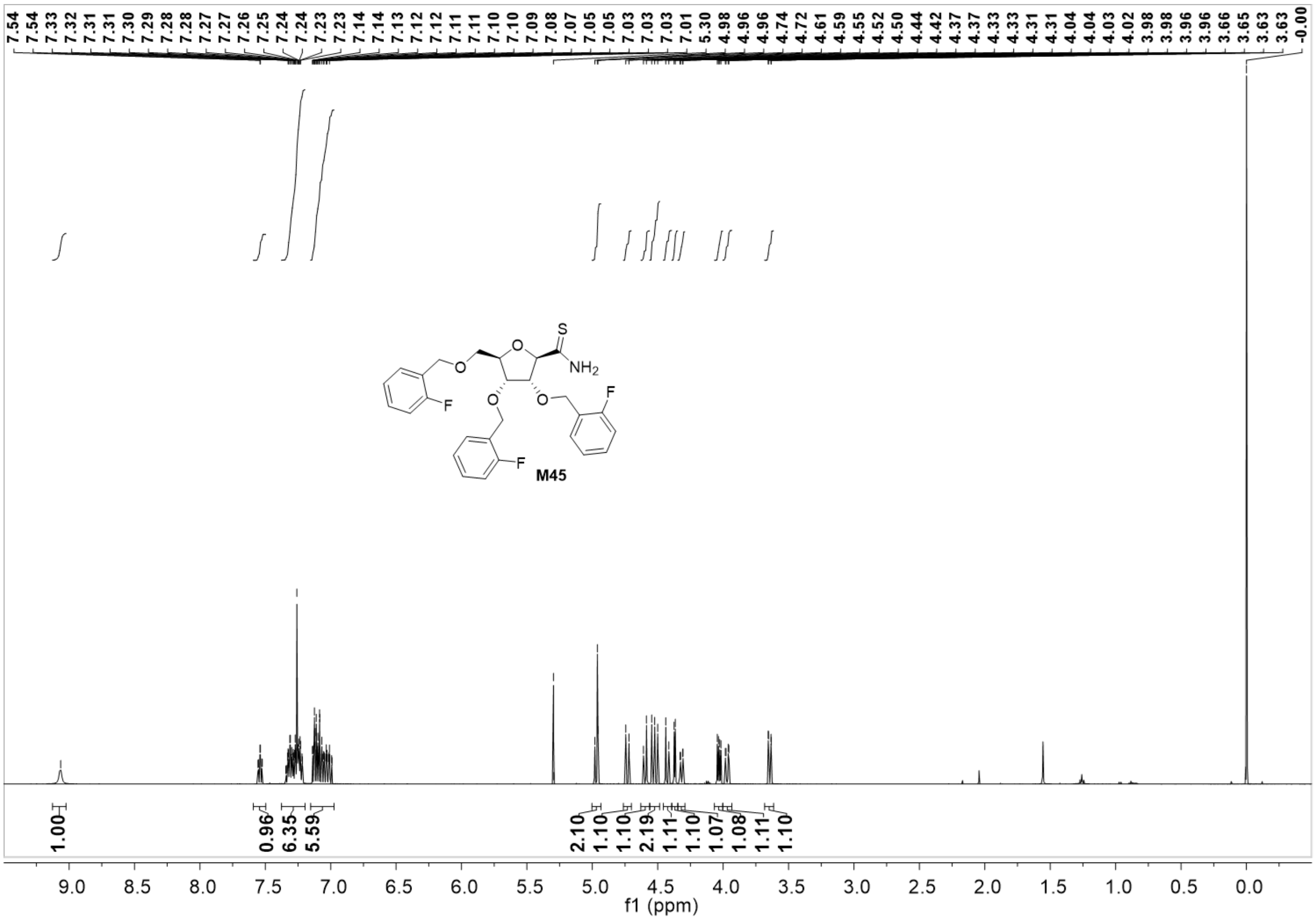
7.54 7.54 7.33 7.32 7.31 7.31 7.30 7.29 7.28 7.28 7.27 7.27 7.26 7.25 7.24 7.24 7.23 7.23 7.14 7.14 7.13 7.12 7.12 7.11 7.11 7.10 7.10 7.09 7.08 7.07 7.05 7.05 7.03 7.03 7.03 7.01 5.30 4.98 4.96 4.96 4.74 4.72 4.61 4.59 4.55 4.52 4.50 4.44 4.42 4.37 4.37 4.33 4.33 4.31 4.31 4.04 4.04 4.03 4.02 3.98 3.98 3.96 3.96 3.66 3.65 3.63 3.63 -0.00
S
O
$NH_2$
O
O
O
F
F
F
M45
1.00
0.96
6.35
5.59
2.10
1.10
1.10
2.19
1.11
1.10
1.07
1.08
1.11
1.10
9.0 8.5 8.0 7.5 7.0 6.5 6.0 5.5 5.0 4.5 4.0 3.5 3.0 2.5 2.0 1.5 1.0 0.5 0.0
f1 (ppm)

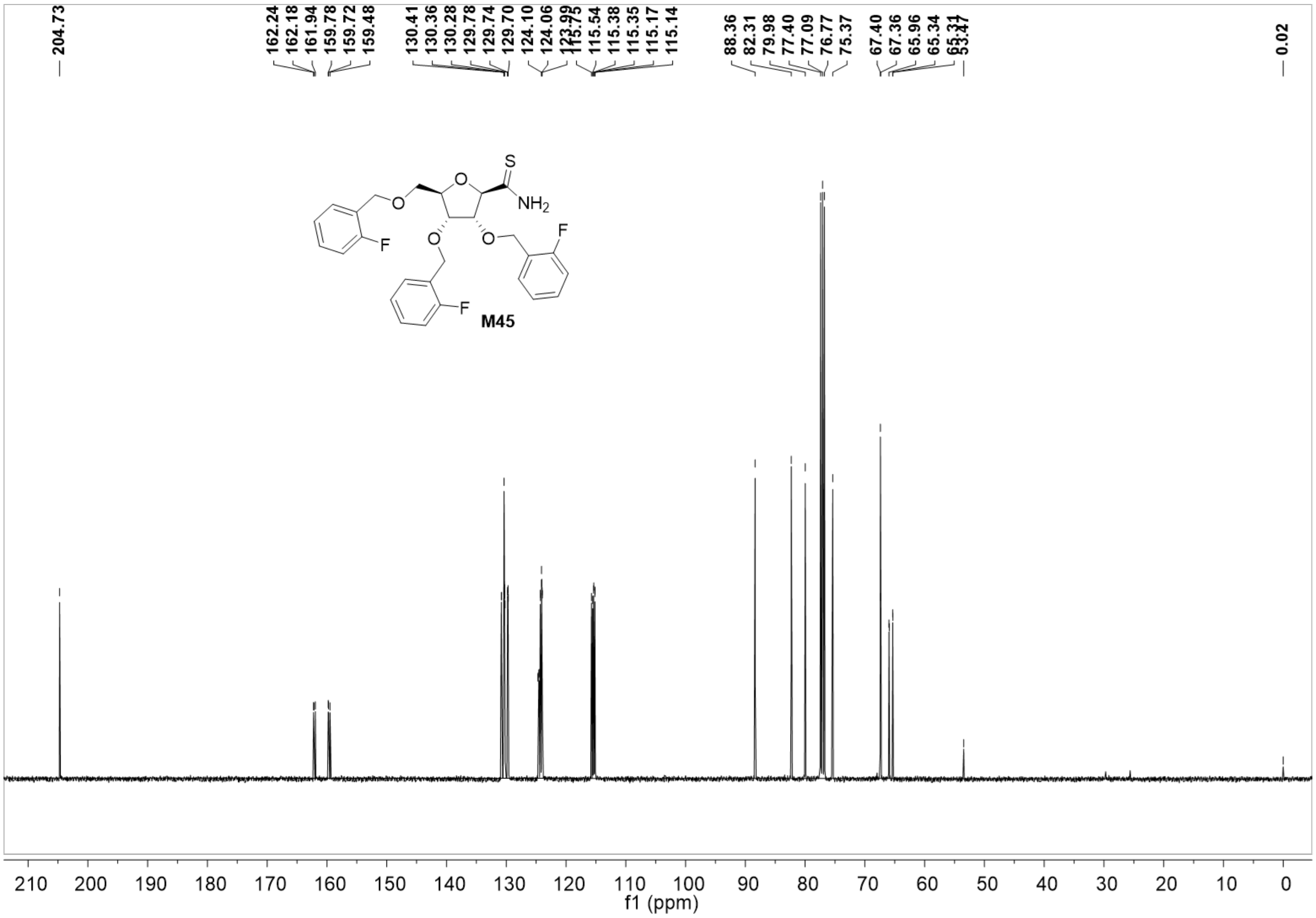

204.73
162.24
162.18
161.94
159.78
159.72
159.48
130.41
130.36
130.28
129.78
129.74
129.70
124.10
124.06
123.99
115.75
115.54
115.38
115.35
115.17
115.14
88.36
82.31
79.98
77.40
77.09
76.77
75.37
67.40
67.36
65.96
65.34
65.31
53.47
0.02
S
NH2
O
F
M45
210 200 190 180 170 160 150 140 130 120 110 100 90 80 70 60 50 40 30 20 10 0
f1 (ppm)

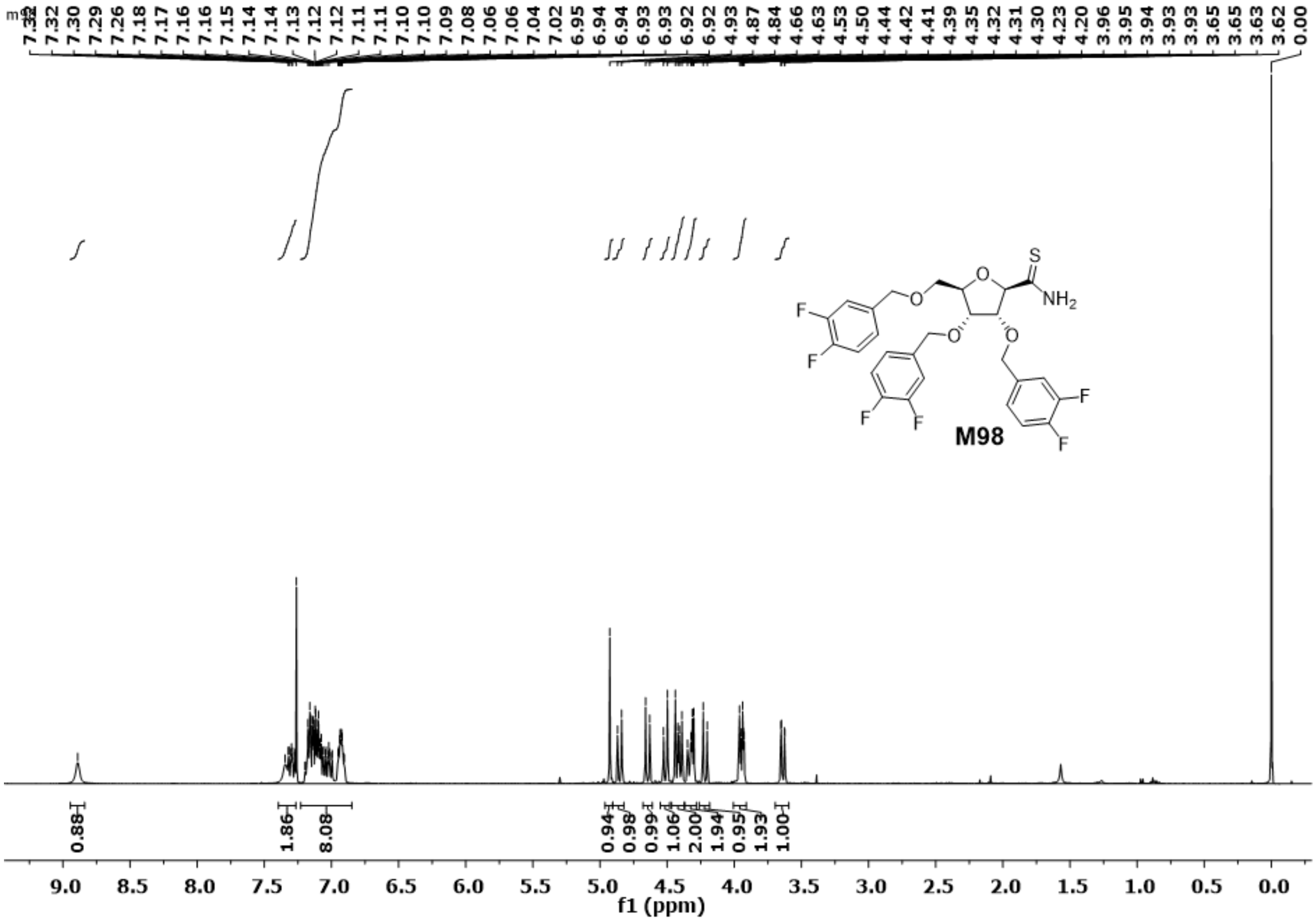

7.32
7.30
7.29
7.26
7.18
7.17
7.16
7.15
7.14
7.14
7.13
7.12
7.12
7.11
7.11
7.10
7.10
7.09
7.08
7.06
7.06
7.04
7.02
6.95
6.94
6.94
6.93
6.93
6.92
6.92
4.93
4.87
4.84
4.66
4.63
4.53
4.50
4.44
4.42
4.41
4.39
4.35
4.32
4.31
4.30
4.23
4.20
3.96
3.95
3.94
3.93
3.93
3.65
3.65
3.63
3.62
0.00
S
NH2
O
F
M98
0.88
1.86
8.08
0.94
0.98
0.99
1.06
2.00
1.94
0.95
1.93
1.00
9.0
8.5
8.0
7.5
7.0
6.5
6.0
5.5
5.0
4.5
4.0
3.5
3.0
2.5
2.0
1.5
1.0
0.5
0.0
f1 (ppm)

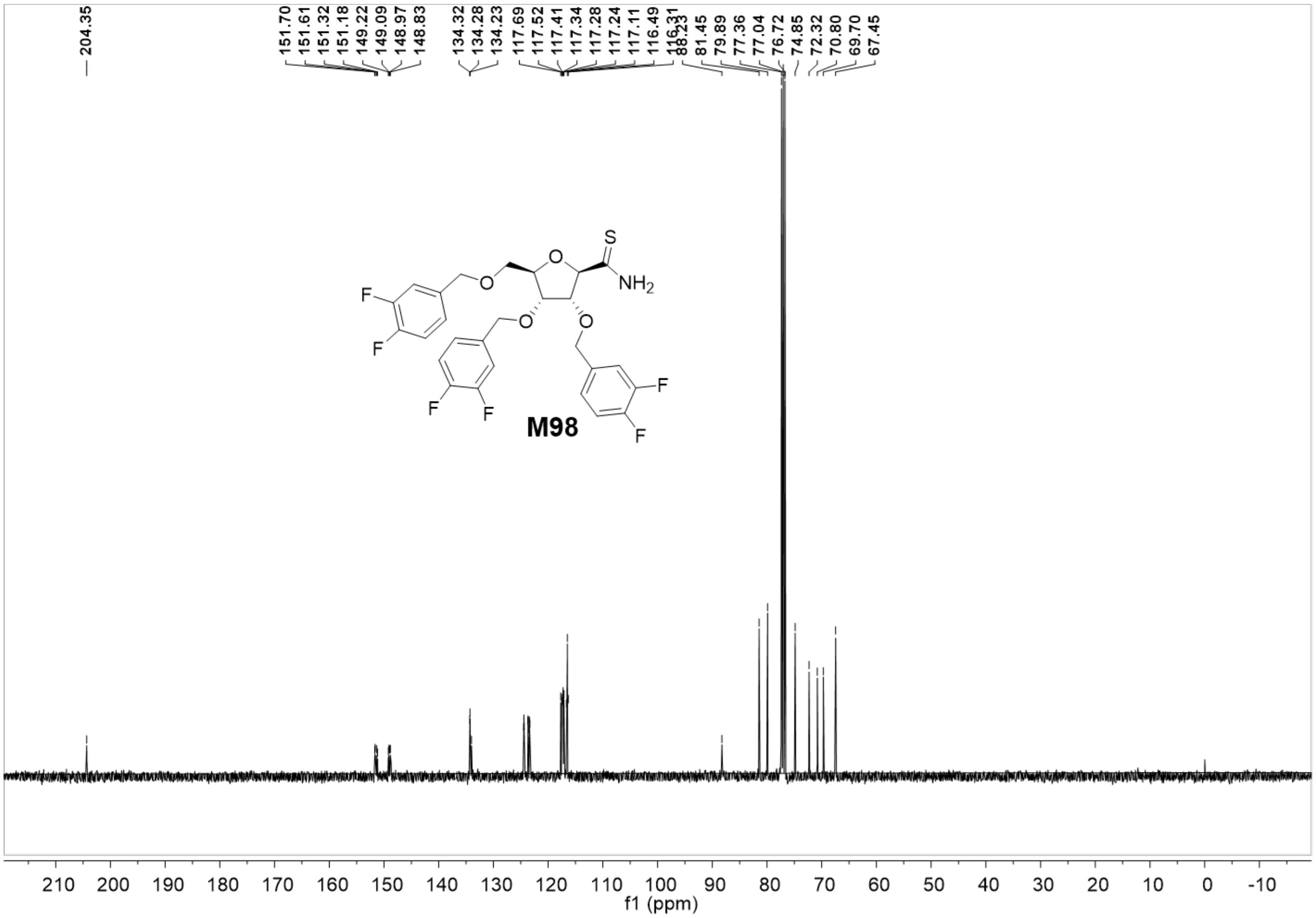
204.35
151.70
151.61
151.32
151.18
149.22
149.09
148.97
148.83
134.32
134.28
134.23
117.69
117.52
117.41
117.34
117.28
117.24
117.11
116.49
116.31
88.23
81.45
79.89
77.36
77.04
76.72
74.85
72.32
70.80
69.70
67.45
S
NH2
O
F
M98
210 200 190 180 170 160 150 140 130 120 110 100 90 80 70 60 50 40 30 20 10 0 -10
f1 (ppm)

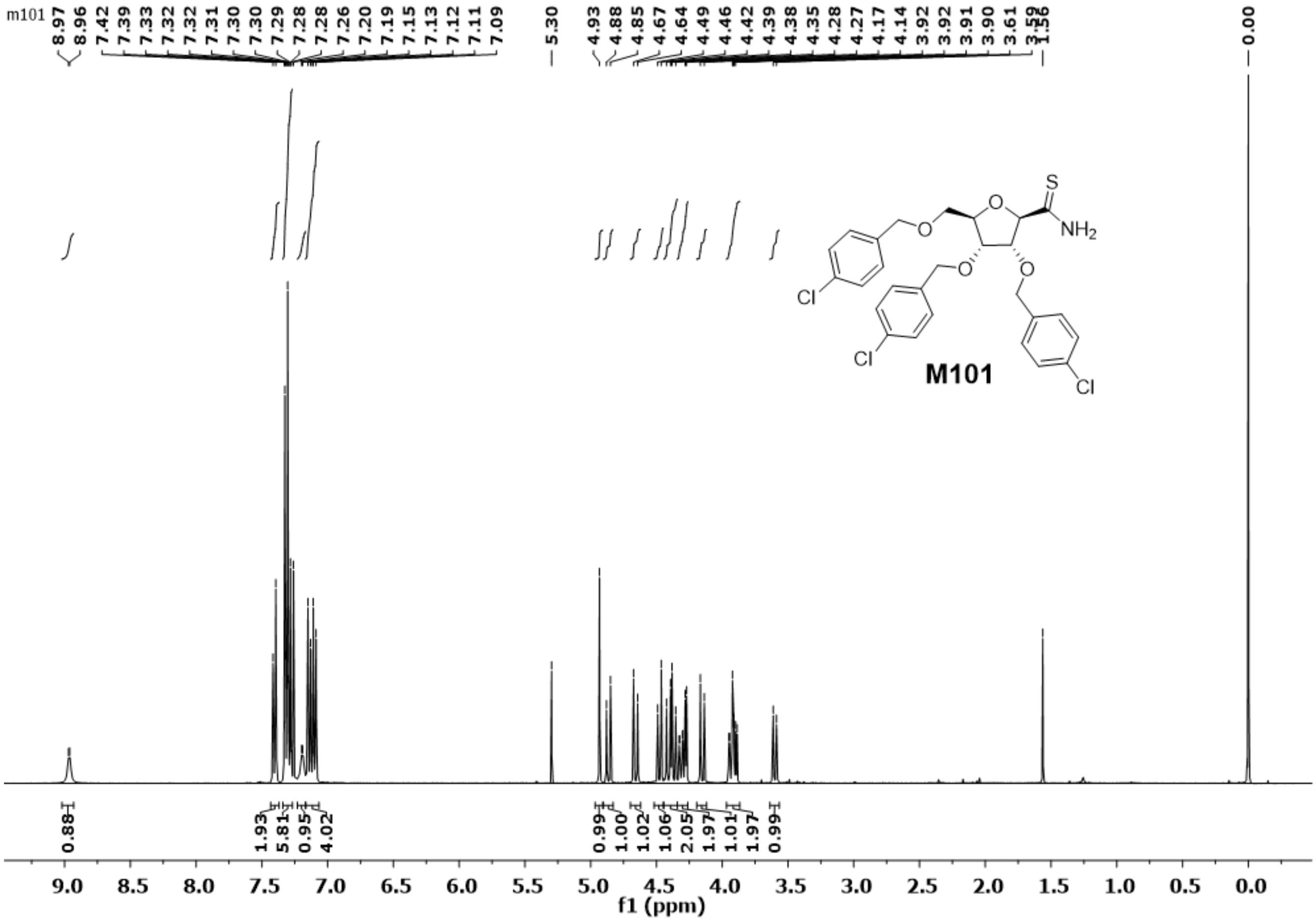
m101
8.97
8.96
7.42
7.39
7.33
7.32
7.32
7.31
7.30
7.29
7.28
7.28
7.26
7.20
7.19
7.15
7.13
7.12
7.11
7.09
5.30
4.93
4.88
4.85
4.67
4.64
4.49
4.46
4.42
4.39
4.38
4.35
4.28
4.27
4.17
4.14
3.92
3.92
3.91
3.90
3.61
3.59
1.56
0.00
S
O
NH2
O
O
O
Cl
Cl
Cl
M101
0.88
1.93
5.81
0.95
4.02
0.99
1.00
1.02
1.06
2.05
1.97
1.01
1.97
0.99
9.0
8.5
8.0
7.5
7.0
6.5
6.0
5.5
5.0
4.5
4.0
3.5
3.0
2.5
2.0
1.5
1.0
0.5
0.0
f1 (ppm)

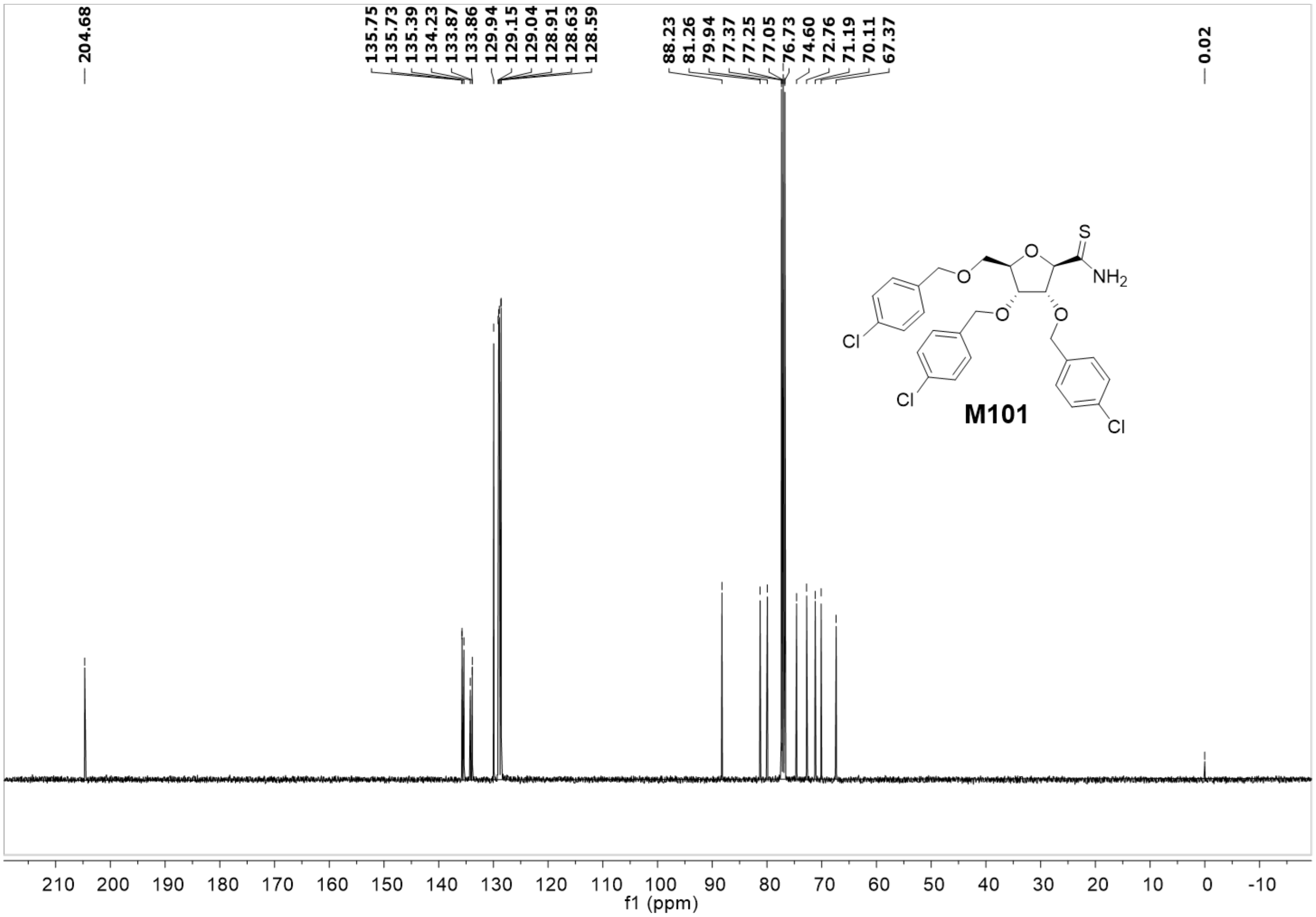

204.68
135.75
135.73
135.39
134.23
133.87
133.86
129.94
129.15
129.04
128.91
128.63
128.59
88.23
81.26
79.94
77.37
77.25
77.05
76.73
74.60
72.76
71.19
70.11
67.37
0.02
S
O
NH2
O
O
O
Cl
Cl
Cl
M101
210 200 190 180 170 160 150 140 130 120 110 100 90 80 70 60 50 40 30 20 10 0 -10
f1 (ppm)

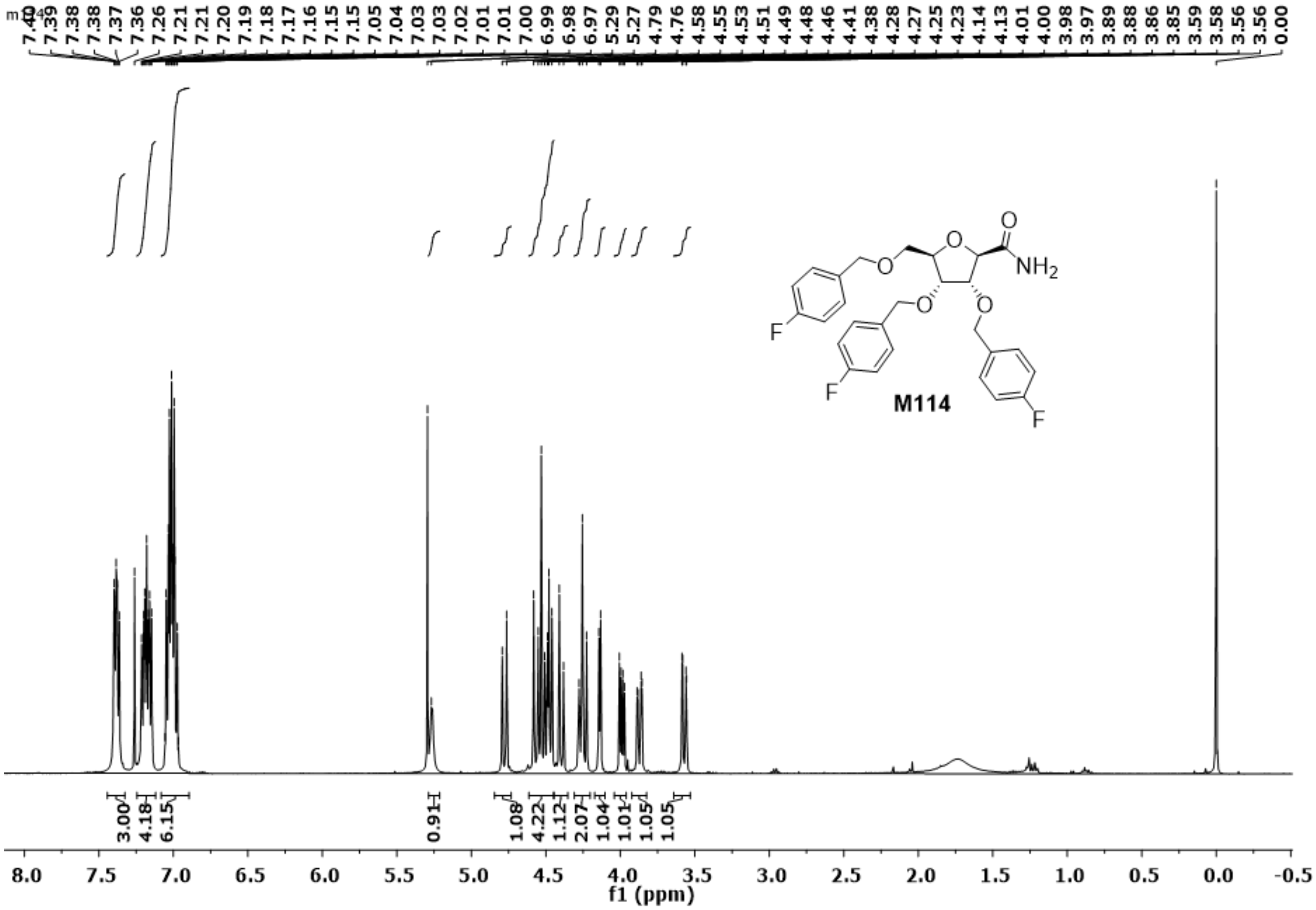
M114
f1 (ppm)
NH2
F
O

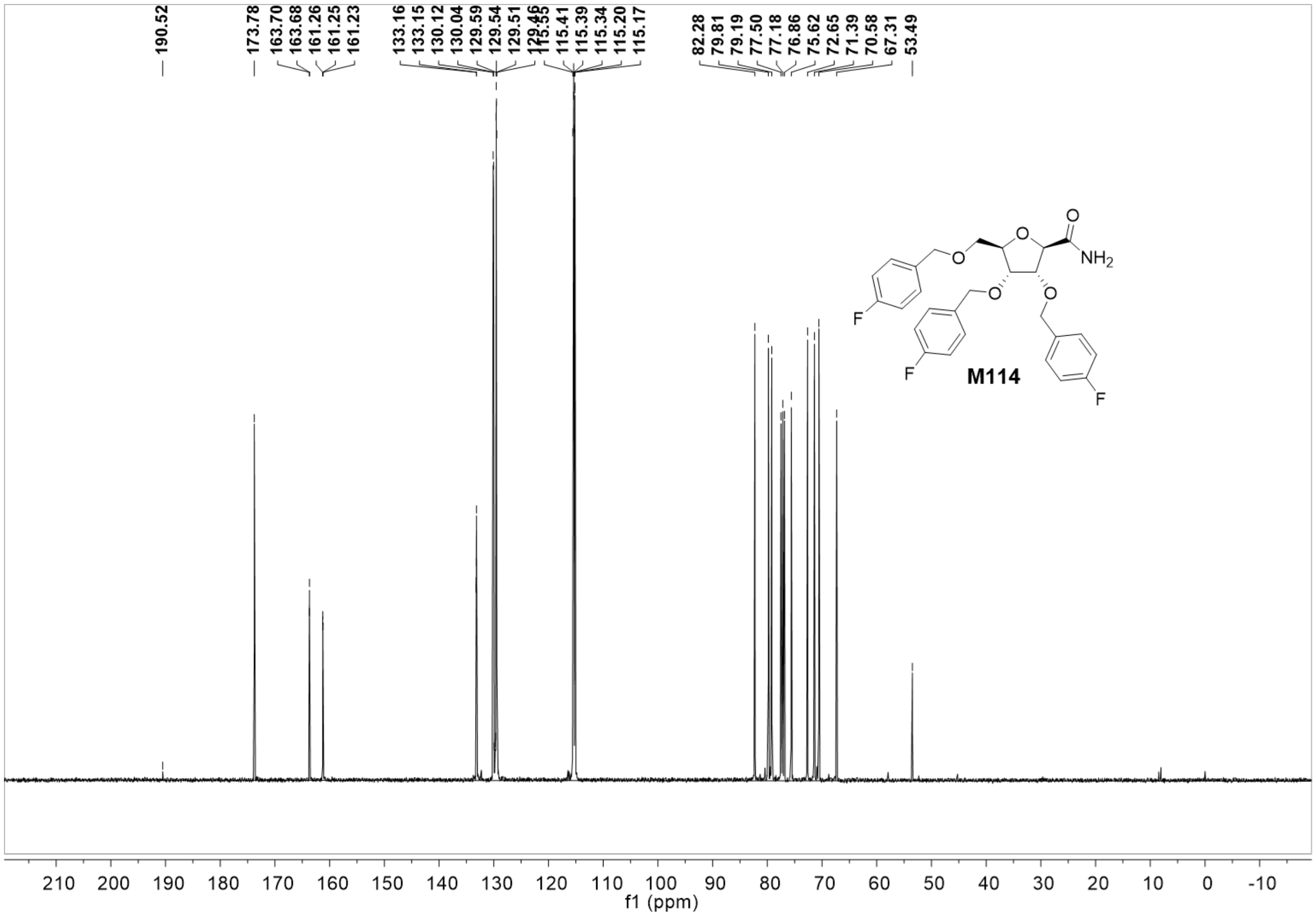
190.52
173.78
163.70
163.68
161.26
161.25
161.23
133.16
133.15
130.12
130.04
129.59
129.54
129.51
129.46
115.55
115.41
115.39
115.34
115.20
115.17
82.28
79.81
79.19
77.50
77.18
76.86
75.62
72.65
71.39
70.58
67.31
53.49
O
NH2
O
O
O
O
F
F
F
M114
210 200 190 180 170 160 150 140 130 120 110 100 90 80 70 60 50 40 30 20 10 0 -10
f1 (ppm)
